# Centrioles control the capacity, but not the specificity, of cytotoxic T cell killing

**DOI:** 10.1101/710053

**Authors:** Fella Tamzalit, Diana Tran, Weiyang Jin, Vitaly Boyko, Hisham Bazzi, Ariella Kepecs, Lance C. Kam, Kathryn V. Anderson, Morgan Huse

## Abstract

Immunological synapse formation between cytotoxic T lymphocytes (CTLs) and the target cells they aim to destroy is accompanied by reorientation of the CTL centrosome to a position beneath the synaptic membrane. Centrosome polarization is thought to enhance the potency and specificity of killing by driving lytic granule fusion at the synapse and thereby the release of perforin and granzymes toward the target cell. To test this model, we employed a genetic strategy to delete centrioles, the core structural components of the centrosome. Centriole deletion altered microtubule architecture, as expected, but surprisingly had no effect on lytic granule polarization and directional secretion. Nevertheless, CTLs lacking centrioles did display substantially reduced killing potential, which was associated with defects in both lytic granule biogenesis and synaptic actin remodeling. These results reveal an unexpected role for the intact centrosome in controlling the capacity, but not the specificity, of cytotoxic killing.

## Introduction

Cytotoxic T lymphocytes (CTLs) and natural killer cells play critical roles in anti-viral and anti-cancer responses by selectively destroying infected or transformed target cells. The most prevalent mechanism of target cell killing involves the secretion of a cellular venom containing the hydrophobic protein perforin and several granzyme proteases (Dustin and Long, 2010; Stinchcombe and Griffiths, 2007). Perforin forms oligomeric pores on the target cell surface, inducing a membrane damage response that allows granzymes to access the cytoplasm, where they cleave specific substrates to induce apoptosis (Thiery and Lieberman, 2014). Essentially all cell types are sensitive to this mode of killing. Hence, specialized mechanisms have evolved to ensure that the effects of perforin and granzyme are constrained to the target cell alone.

Cytotoxic lymphocytes store perforin and granzyme in specialized secretory lysosomes called lytic granules (Stinchcombe and Griffiths, 2007; Thiery and Lieberman, 2014), whose low pH environment quenches the activity of both proteins. Within minutes of target cell recognition, these granules traffic along microtubules to the stereotyped interface between the lymphocyte and the target cell, which is known as the immunological synapse (IS). Here, they fuse with the plasma membrane to release their contents into the intercellular space. Concomitantly, cortical F-actin at the IS undergoes dramatic remodeling, generating a complex landscape of highly dynamic sheets and protrusions. Recent studies indicate that these structures boost the lytic activity of perforin by applying mechanical force against the target cell (Basu et al., 2016; Tamzalit et al., 2019).

Effective killing depends on the exclusive release of granule contents at the IS. This concentrates the perforin and granzymes delivered to the target while simultaneously minimizing damage to bystander cells in the surrounding tissue. Directional granule release (also called degranulation) is generally thought to depend on the centrosome, a membraneless organelle that serves as the microtubule-organizing center (MTOC) in most cells (Bettencourt-Dias and Glover, 2007; Pihan, 2013). Microtubules radiate from the centrosome with their minus ends directed inward and the their plus ends outward. Metazoan centrosomes contain at least two centrioles, which are cylindrical structures made of parallel microtubule triplets, surrounded by a cloud of proteinaceous pericentriolar material (PCM). The centrioles maintain centrosomal organization, while the PCM, which is highly enriched in γ-tubulin, is responsible for nucleating microtubule growth. A defining feature of IS formation is the movement of the centrosome to a position just beneath the center of the interface. Lytic granules cluster around the centrosome in activated lymphocytes, and therefore its reorientation to the IS positions the granules close to the synaptic membrane (Liu and Huse, 2015; Stinchcombe and Griffiths, 2007). The temporal coordination between granule clustering, centrosome reorientation, and target cell killing implies that the centrosome delivers the granules to the IS for fusion, thereby maintaining the potency and specificity of the response. Consistent with this model, multiple studies have associated delayed or impaired centrosome reorientation with reduced cytotoxicity (de la Roche et al., 2013; Jenkins et al., 2014; Quann et al., 2009; Tsun et al., 2011). Furthermore, deletion of Cep83, a centrosomal protein required for centriole docking with the plasma membrane, was found to inhibit CTL degranulation (Stinchcombe et al., 2015). Other studies, however, have suggested that robust CTL- and NK cell-mediated killing occurs in the absence of centrosome polarization (Bertrand et al., 2013; Butler and Cooper, 2009; Chauveau et al., 2010). Hence, the precise role of the centrosome during cytotoxic responses remains unsettled.

In the present study, we employed a genetic strategy to delete centrioles from CTLs and thereby investigate the importance of the centrosome for CTL function. Centriole deletion markedly reduced cytotoxicity, as expected. This defect, however, did not result from impaired directional secretion. Indeed, lytic granule trafficking and release remained polarized toward the IS. Rather, centriole deficient CTLs exhibited dysregulated granule biogenesis, which diminished their steady state perforin and granzyme stores and limited their cytotoxic potential. Centriole loss also impaired synaptic F-actin remodeling, reducing force exertion across the IS. These results demonstrate that the centrosome controls the capacity, but not the specificity, of target cell killing and reveal an unexpected link between microtubule and F-actin dynamics during T cell activation.

## Results

### *Sas4*^*−/−*^ *Trp53*^*−/−*^ CTLs lack centrioles and exhibit perturbed centrosome and microtubule architecture

The centriolar adaptor protein spindle assembly defective-4 (SAS4, also called CENPJ) is required for the formation and maintenance of centrosomes in eukaryotes (Gopalakrishnan et al., 2011; Kirkham et al., 2003). In the absence of SAS4, most cell types undergo P53 dependent apoptosis downstream of a mitotic centriole checkpoint (Bazzi and Anderson, 2014). Cells lacking both SAS4 and P53, however, are healthy in culture and proliferate at only a slightly slower rate than normal. Hence, to generate CTLs lacking centrioles, we crossed mice bearing conditional “floxed” alleles of *Sas4* and *Trp53* (the P53 gene) with transgenic mice expressing the OT1 TCR, which recognizes the ovalbumin_257-264_ peptide presented by the class I MHC protein H2-K^b^ (H2-K^b^-OVA). Lymphocytes from OT1-*Sas4*^fl/fl^*Trp53*^fl/fl^ animals were activated in vitro with their cognate antigen and then retrovirally transduced 48 hours later to express Cre recombinase (Fig. 1A). After 5-6 days of additional culturing in the presence of IL2 and FACS sorting for transduced cells, differentiated OT1-*Sas4*^−/−^*Trp53*^−/−^ (DKO) CTLs were obtained in sufficient numbers for imaging experiments and functional studies (Fig. S1A).

**Figure 1.**
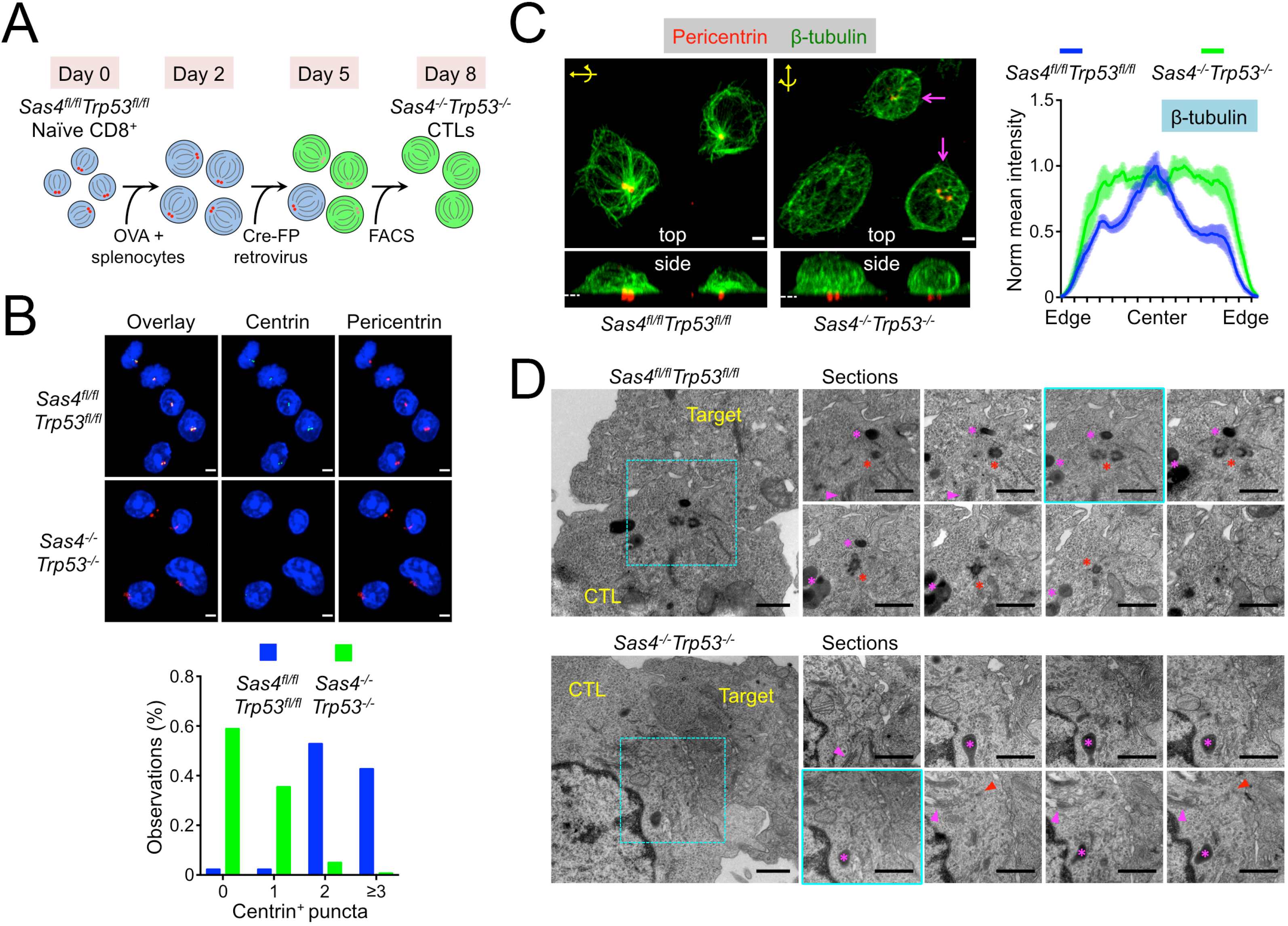
*Sas4*^*−/−*^*Trp53*^*−/−*^ CTLs lack centrioles and display altered centrosomal and microtubule architecture. (A) Schematic diagram of protocol for generating OT1 *Sas4^−/−^ Trp53^−/−^* CTLs. FP indicates GFP, CFP, or RFP. (B) Above, representative confocal images of *Sas4*^*fl/fl*^*Trp53*^*fl/fl*^ and *Sas4*^*−/−*^*Trp53*^*−/−*^ OT1 CTLs stained with antibodies against the indicated centriolar and centrosomal proteins. Nuclear DAPI staining is shown in blue. Scale bars = 3 μm. Below, quantification of centrin^+^ puncta in *Sas4*^*fl/fl*^*Trp53*^*fl/fl*^ (N = 89) and *Sas4*^*−/−*^*Trp53*^*−/−*^ (N = 142) OT1 CTLs. (C) Left, representative confocal images of *Sas4*^*fl/fl*^*Trp53*^*fl/fl*^ and *Sas4*^*−/−*^*Trp53*^*−/−*^ OT1 CTLs stained with antibodies against pericentrin and β-tubulin. Top view (z-projection) images are shown above with corresponding side views (x- or y-projections) below. Dotted white lines indicate the plane of the IS in the side views. The axis and rotation used to generate the side view is indicated in yellow in the top views. Magenta arrows indicate CTLs with residual centralized microtubule organization. Scale bars = 2 μm. Right, normalized, mean β-tubulin fluorescence intensity in linescans from one cell edge to the opposite edge (see Fig. S1C and Methods, N ≥ 12 for each cell type). Error bars denote standard error of the mean (SEM). (D) TEM analysis of conjugates formed between *Sas4*^*fl/fl*^*Trp53*^*fl/fl*^ or *Sas4*^*−/−*^*Trp53*^*−/−*^ OT1 CTLs and OVA-loaded EL4 target cells. Wide view panels of representative conjugates are shown on the left, with serial sections of the central synaptic domain (indicated by the dashed cyan box in each wide view image) shown to the right. The section corresponding to the wide view image is outlined in cyan. Red asterisks indicate centrioles in the *Sas4*^*fl/fl*^*Trp53*^*fl/fl*^ CTL and red arrowheads denote residual clusters of electron dense, PCM-like material in the *Sas4*^*−/−*^*Trp53*^*−/−*^ CTL. Magenta asterisks and arrowheads indicate lytic granules Golgi stacks, respectively. Scale bars = 1 μm. All data are representative of at least two independent experiments.

Immunocytochemical analysis using antibodies against centrin, a centriole marker, revealed that a majority of DKO CTLs lacked any detectable centrioles, with most of the remaining cells containing only one centrin^+^ object (Fig. 1B). This phenotype was readily apparent by day 6 (four days after Cre transduction), and became even more pronounced thereafter (Fig. S1B). By contrast, most wild type (OT1-*Sas4*^fl/fl^*Trp53*^fl/fl^) CTLs contained at least two detectable centrioles at all time points examined (Fig. 1B and S1B). This is typical for normal cells, which contain between two and four centrioles, depending on the stage of the centrosome cycle (Bettencourt-Dias and Glover, 2007). Wild type CTLs also exhibited focal accumulation of the PCM proteins pericentrin and γ-tubulin, which tightly colocalized with centrin^+^ objects. Pericentrin and γ-tubulin staining was less organized in DKO CTLs, often appearing as a smattering of puncta rather than two intensely fluorescent foci (Fig. 1B, S1C). Nevertheless, both markers remained clustered on one side of the cell, implying some degree of residual PCM organization in the absence of centrioles. Importantly, *Trp53*^−/−^ CTLs with one functional copy of *Sas4* displayed normal centrosomal architecture (Fig. S2A), indicating that the DKO phenotype resulted from SAS4 and not P53 deficiency.

Next, we examined microtubule organization by staining OT1 CTLs on stimulatory coverslips coated with H2-K^b^-OVA and ICAM1, a ligand for the α_L_β_2_ integrin LFA1. OT1 CTLs form IS like contacts with these surfaces, complete with centrosome reorientation toward the interface. In wild type cells, the dense cluster of microtubules radiating from the centrosome was easily observed in z-projection (top view) images (Fig. 1C). This cluster was less obvious in DKO CTLs; indeed, quantification of microtubule density as a function of radial position within cells revealed a marked reduction of centralized microtubule organization (Fig. 1C, Fig. S1D). DKO CTLs did retain some degree of this centralized configuration (magenta arrows in Fig. 1C), however, possibly reflecting the persistence of polarized PCM in the space formerly occupied by the centrosome. We also found that centriole loss did not affect the focal architecture of the Golgi apparatus (Fig. S1E), consistent with previous work indicating that Golgi organization persists after centrosome ablation (Efimov et al., 2007). Given that PCM and the Golgi can both nucleate microtubules, it is possible that they serve as a rudimentary MTOC in the absence of an intact centrosome. Consistent with this idea, both the residual PCM (detected by pericentrin staining) and the Golgi polarized toward the IS in DKO CTLs (Fig. 1C, S1E), much like the centrosome in wild type cells.

To investigate the architectural consequences of *Sas4* deletion at higher resolution, we imaged conjugates between OVA-loaded EL4 cells and either wild type or DKO OT1 CTLs by transmission electron microscopy (TEM). In wild type CTLs, centriole pairs, which appeared as orthogonally oriented cylinders of microtubules, were routinely found in close apposition with the synaptic membrane (Fig. 1D), in agreement with previous studies (Stinchcombe et al., 2006; Stinchcombe et al., 2015). The cytoplasm surrounding the centrioles tended to be enriched in microtubules, Golgi membranes, and lytic granules, consistent with the centrosome serving as an MTOC. By contrast, many DKO CTLs completely lacked centriole barrels, a phenotype that we confirmed by analyzing serial sections through the entire contact site (Fig. 1D). Interestingly, in some DKO CTLs we observed small accumulations of electron dense material at the IS that appeared to be residual clusters of PCM or centriole fragments (black arrowheads in Fig 1D). Similar structures were observed in a prior study of acentriolar DT40 B cells (Sir et al., 2013). Despite the absence of centrioles, the Golgi stacks remained polarized to the IS in DKO CTLs, consistent with the fluorescence imaging data described above (Fig. S1E). We also observed numerous microtubules in DKO synapses, although it was difficult to determine whether they were organized by the Golgi or by the PCM-like electron dense material. Taken together, these data indicate that DKO CTLs lack centrioles and exhibit altered, but not completely disrupted, microtubule and centrosome organization.

### Centriole deletion does not block TCR signaling

Having confirmed that DKO CTLs lack centrioles, we investigated their responses to TCR stimulation. CTLs were mixed with beads containing immobilized H2-K^b^-OVA and ICAM1, a ligand for the α_L_β_2_ integrin LFA1. In DKO CTLs, this induced degradation of IκB and rapid phosphorylation of Erk1/2 and AKT, indicative of strong signaling through the MAP kinase, phosphoinositide 3-kinase, and NFκB pathways (Fig. 2A). These responses were largely indistinguishable from those of control CTLs, indicating that centriole loss does not grossly impair TCR signaling. We also examined calcium (Ca^2+^) influx downstream of the TCR by imaging CTLs loaded with the Ca^2+^ sensitive dye Fura2 on supported bilayers bearing H2-K^b^-OVA and ICAM1. DKO CTLs responded as well as, if not better than, control cells in these experiments (Fig. 2B), further supporting the conclusion centrioles are not required for T cell activation.

**Figure 2.**
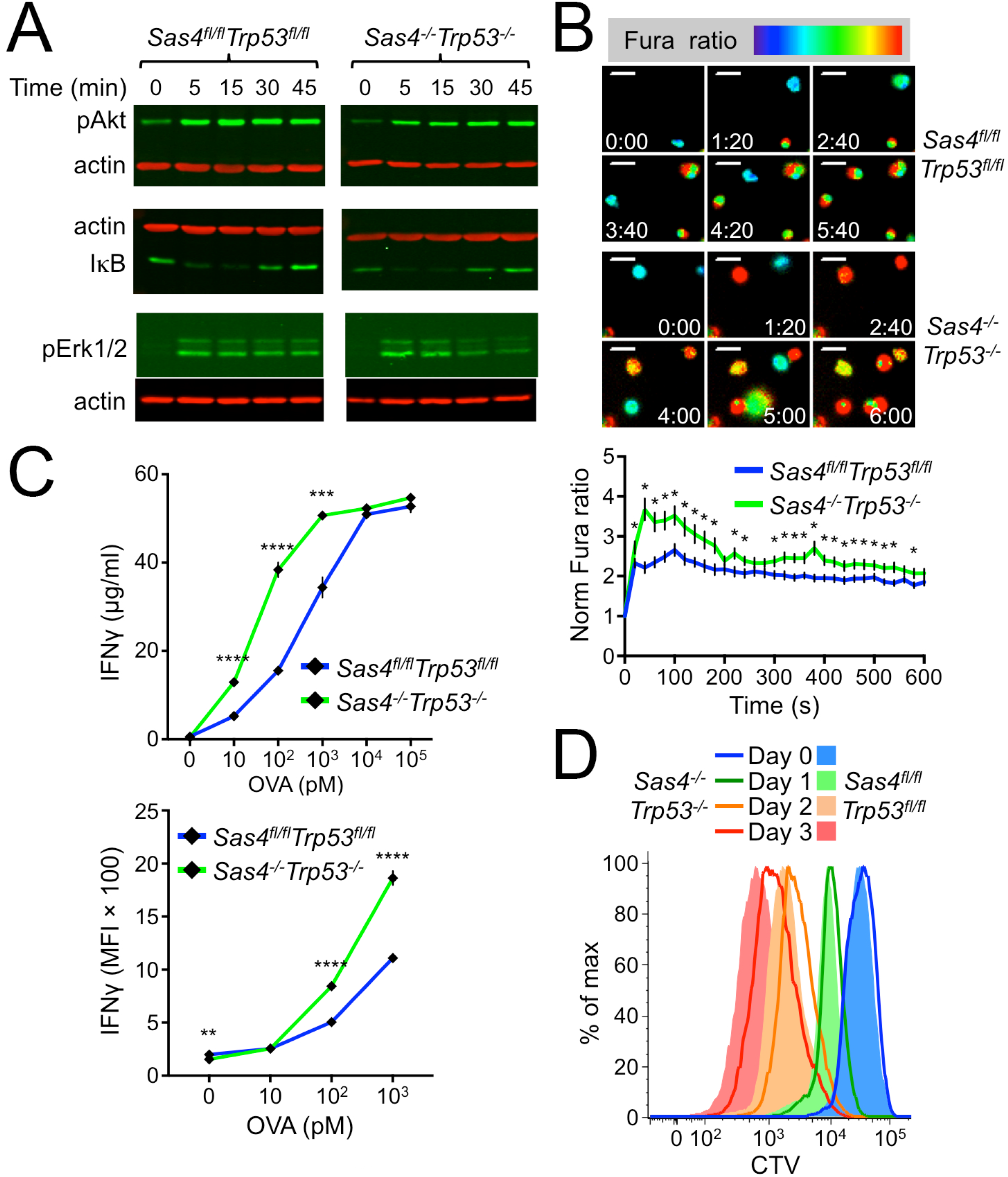
*Sas4*^*−/−*^*Trp53*^*−/−*^ CTLs respond to TCR stimulation. (A) *Sas4*^*fl/fl*^*Trp53*^*fl/fl*^ and *Sas4*^*−/−*^*Trp53*^*−/−*^ OT1 CTLs were mixed with beads coated with H2-K^b^-OVA and ICAM1. At the indicated times, pAKT, IκB, and pErk1/2 were assessed by immunoblot, using actin as a loading control. (B) *Sas4*^*fl/fl*^*Trp53*^*fl/fl*^ and *Sas4*^*−/−*^*Trp53*^*−/−*^ OT1 CTLs were loaded with Fura2-AM and imaged on glass surfaces coated with H2-K^b^-OVA and ICAM1. Above, representative time-lapse montages showing Fura ratio in pseudocolor, with warmer colors indicating higher intracellular Ca^2+^ concentrations. Time in M:SS is shown in each image. Scale bars = 10 μm. Below, mean normalized Fura ratio was graphed against time. N ≥ 29 for each cell type. P values (* indicates P < 0.05) were calculated by two-tailed Student’s T-test. (C) *Sas4*^*fl/fl*^*Trp53*^*fl/fl*^ and *Sas4*^*−/−*^*Trp53*^*−/−*^ OT1 CTLs were mixed with RMA-s cells pulsed with the indicated concentrations of OVA. Antigen-induced IFNγ production was assessed by ELISA (above) and intracellular staining (below). P values (**, ***, and **** indicate P < 0.01, P < 0.001, and P < 0.0001, respectively) were calculated by two-tailed Student’s T-test. (D) *Sas4*^*fl/fl*^*Trp53*^*fl/fl*^ and *Sas4*^*−/−*^*Trp53*^*−/−*^ OT1 CTLs were incubated with OVA-loaded C57BL/6 splenocytes, and proliferation assessed by CTV dilution at the indicated time points. In B and C, error bars denote SEM. All data are representative of at least two independent experiments.

Next, we assessed the effects of *Sas4* deletion on the secretion of the inflammatory cytokine interferon-γ (IFNγ), a well-established downstream response to TCR stimulation. Co-incubation of OT1 CTLs with antigen-loaded RMA-s target cells induced robust IFNγ production, which could be detected both by intracellular staining and by ELISA. DKO CTLs consistently generated more IFNγ than their wild type counterparts (Fig. 2C). Although we cannot explain this surprising increase, it is possible that it is related to their enhanced Ca^2+^ responses (Fig. 2B). We also examined TCR-induced cell division by stimulating wild type and DKO CTLs with OVA-loaded splenocytes. Although both cell types exhibited strong proliferation in these experiments, cell division was somewhat delayed in DKO cells relative to controls, a difference that was most obvious 48 and 72 hours after stimulation (Fig. 2D). This phenotype was consistent with previous work indicating that centriole loss extends the time spent in mitosis (Bazzi and Anderson, 2014).

Finally, we examined cell surface levels of the TCR and LFA1. TCR expression was consistently higher in DKO CTLs than in wild type controls (Fig. S3A), which potentially explains their enhanced Ca^2+^ and cytokine responses. DKO CTLs exhibited normal ligand-induced TCR downregulation, however, implying that receptor endocytosis in unaffected by centriole loss (Fig. S3B). LFA1 expression, for its part, was identical in both groups of CTLs (Fig. S3A). Collectively, these data indicate that, despite lacking centrioles, DKO CTLs express key activating receptors and respond robustly to antigenic stimulation.

### Centriole deletion reduces cytotoxic capacity but not cytotoxic specificity

To assess the importance of centrioles for cytotoxicity, OT1 CTLs were mixed with RMA-s cells that had been loaded with increasing amounts of OVA (Fig. 3A). DKO CTLs displayed a dramatic cytotoxicity defect at low effector to target (E:T) ratios (1:10, Fig. 3B). At higher E:T (1:1), however, this phenotype was much less apparent (Fig. 3B), and even absent in certain experiments. Importantly, OT1-*Sas4*^+/−^*Trp53*^−/−^ CTLs killed normally at both E:T ratios (Fig. S2B), implying that the DKO defect resulted from SAS4, rather than P53, deficiency. We also measured cytotoxicity at a single cell level by imaging CTLs and target cells that had been loaded into 50 μm × 50 μm polydimethylsiloxane microwells. This approach prevents the formation of large cell clusters, enabling observation of individual cytotoxic interactions for an extended period of time (10-12 hours) (Le Floc'h et al., 2013). Target cell killing was visualized using propidium iodide (PI), a fluorescent DNA intercalating agent, which stains dying cells with compromised plasma membrane integrity (Fig. S4A). The results of these experiments mirrored those of the bulk assays. In microwells containing only one CTL and 1-3 target cells, DKO CTLs exhibited a marked killing defect relative to wild type controls (Fig. S4B-C). However, in microwells containing two CTLs and one target, the likelihood of target cell killing was nearly identical. The fact that DKO cytotoxicity was rescued, in both bulk and single cell experiments, by adding more CTLs suggests that these cells can efficiently kill a small number of targets, but that their potential for extensive serial killing is limited. In other words, our results imply a link between the centriole and CTL killing capacity.

**Figure 3.**
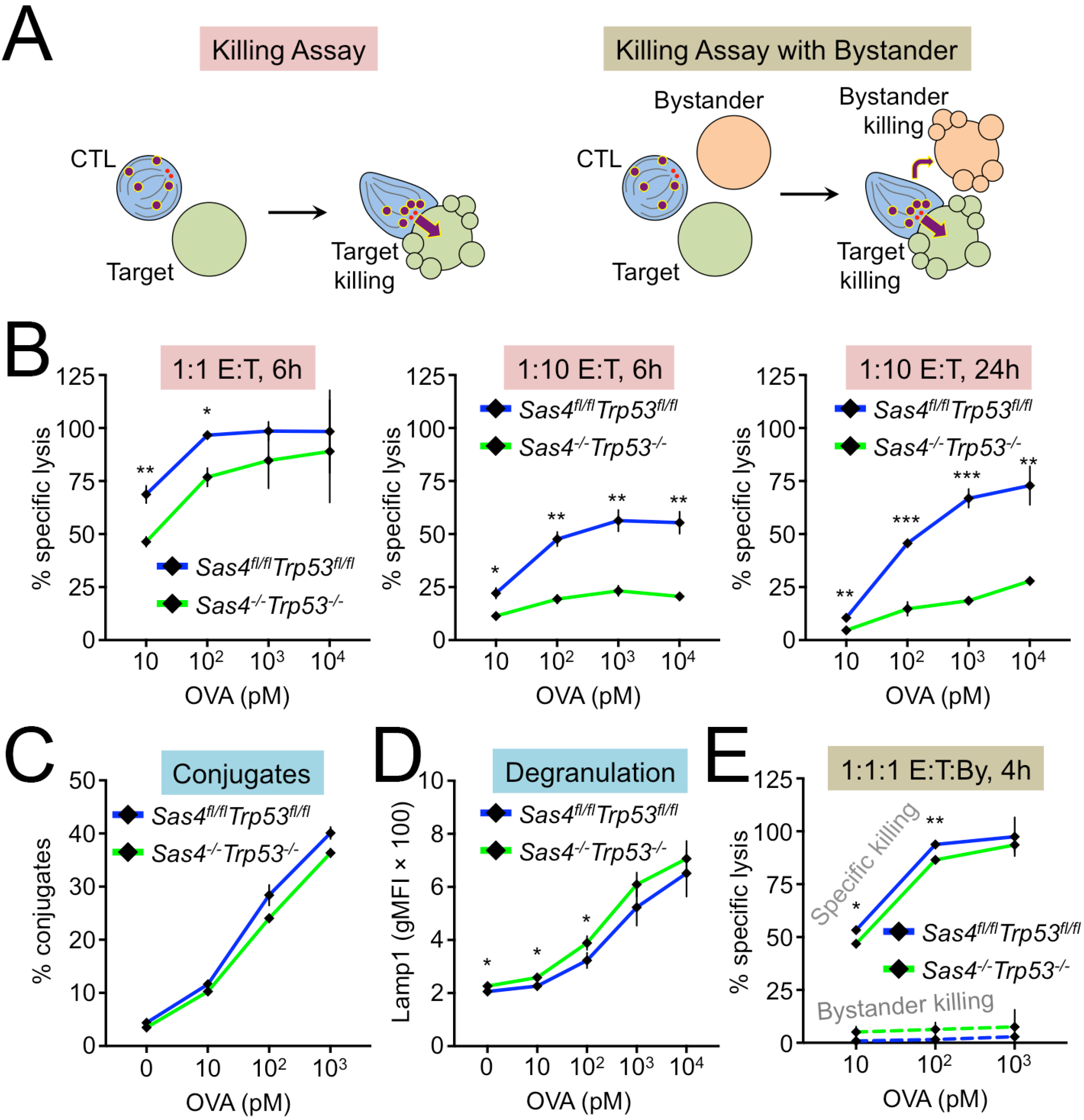
Centriole loss affects killing capacity but not killing specificity. (A) Experimental design for killing assays with (right) and without (left) bystander cells. (B-D) RMA-s target cells were loaded with increasing concentrations of OVA and mixed with *Sas4*^*fl/fl*^*Trp53*^*fl/fl*^ or *Sas4*^*−/−*^*Trp53*^*−/−*^ OT1 CTLs. Specific lysis of RMA-s cells was assessed at the indicated E:T ratios and times. (C) CTL-target cell conjugate formation measured by flow cytometry. (D) Degranulation measured by surface exposure of Lamp1. (E) RMA-s target cells were loaded with increasing concentrations of OVA and mixed with unpulsed RMA-s cells (bystanders) and either *Sas4*^*fl/fl*^*Trp53*^*fl/fl*^ or *Sas4*^*−/−*^*Trp53*^*−/−*^ OT1 CTLs at a 1:1:1 ratio. Specific lysis of both targets and bystanders was assessed after 4h. P values (*, **, and ***, indicate P < 0.05, P < 0.01, and P < 0.001, respectively) were calculated by two-tailed Student’s T-test in all panels. Error bars denote SEM. All data are representative of at least two independent experiments.

Effective target cell killing requires that CTLs conjugate tightly with target cells and then degranulate. Conjugate formation, which we measured using a flow cytometry-based approach, was normal in DKO CTLs (Fig. 3C). To quantify degranulation, we monitored cell surface exposure of the lysosomal protein Lamp1. TCR stimulation of DKO CTLs induced robust surface expression of Lamp1 that was essentially indistinguishable from the response in wild type cells (Fig. 3D). Hence, the reduced killing capacity of DKO CTLs did not result from impaired conjugate formation or lytic granule release.

Given the purported role of the centrosome in mediating granule polarization to the IS, we hypothesized that centriole loss impairs target cell killing by disrupting the directional secretion of cytotoxic factors. Unpolarized release of perforin and granzyme would be expected to have two functional consequences, 1) reducing the killing of antigen bearing target cells, and 2) increasing the killing of bystander cells. To test the second prediction, we performed cytotoxicity assays in which CTLs were mixed with both OVA-loaded (target) and unloaded (bystander) RMA-s cells at a 1:1:1 ratio (Figure 3A). Surprisingly, DKO CTLs induced little to no bystander cell killing, despite driving robust destruction of bone fide targets (Figure 3E). We observed nearly identical results with wild type CTLs, implying that both populations kill with equivalent selectivity. Taken together, these results suggest that the CTL centriole controls the capacity, but not the specificity, of cytotoxic responses.

### Microtubules are dispensable for directional release of cytotoxic proteins

The observation that centriole loss did not affect the specificity of killing called into question whether an intact centrosome is actually necessary for polarized lytic granule release at the IS. To investigate this issue, we imaged CTLs that expressed a degranulation reporter comprising a pH sensitive GFP (pHluorin) fused to the C-terminal tail of Lamp1 (Rak et al., 2011). pHluorin-Lamp1 accumulates in lytic granules, where its fluorescence is quenched by the low pH environment. Granule fusion with the plasma membrane, however, neutralizes the pH around the pHluorin, allowing it to fluoresce (Fig. 4A). Degranulating CTLs were imaged on stimulatory glass surfaces coated with H2-K^b^-OVA and ICAM1 (Fig. S5A). Degranulation events manifested as abrupt increases in green fluorescence, which often appeared and then disappeared in consecutive time points (Fig. 4B). In wild type cells, degranulations clustered close to the glass surface (within ~ 2 μm), indicative of synaptic exocytosis (Fig. 4C). Remarkably, OT1-DKO CTLs also exhibited highly polarized granule release, implying that centriole loss does not compromise directional secretion.

**Figure 4.**
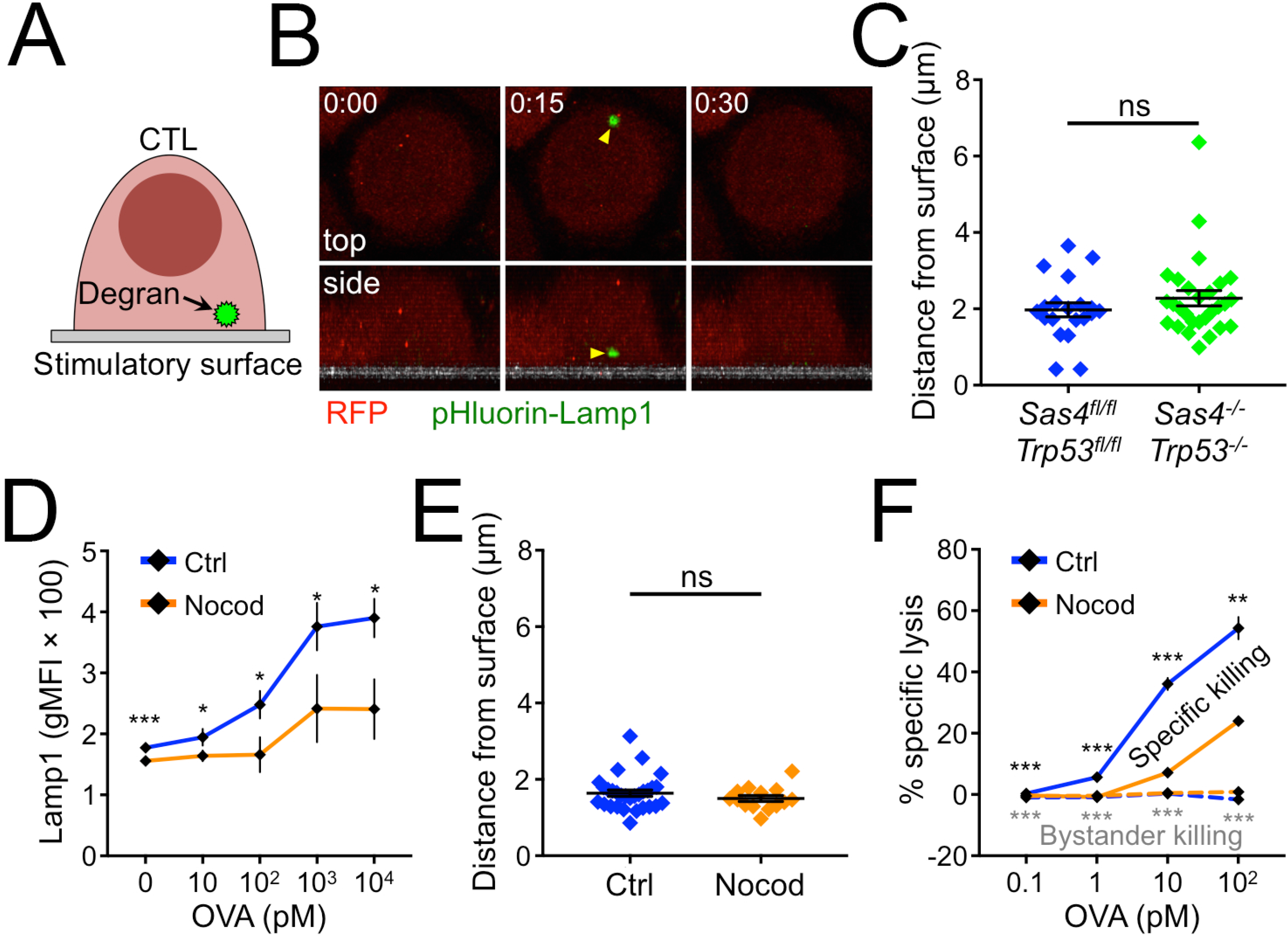
*Sas4*^*−/−*^*Trp53*^*−/−*^ CTLs degranulate at the IS. (A) Schematic diagram showing pHluorin-Lamp1-based detection of degranulation on stimulatory surfaces. (B-C) *Sas4*^*fl/fl*^*Trp53*^*fl/fl*^ or *Sas4*^*−/−*^*Trp53*^*−/−*^ OT1 CTLs expressing RFP and pHluorin-Lamp1 were imaged by confocal microscopy on glass surfaces coated with H2-K^b^-OVA and ICAM1. A time-lapse montage of a representative degranulation event (yellow arrowhead), with top views shown above and side views below. The surface is indicated in gray in the side views. Time in M:SS is shown in the top left corner of each top view image. (C) Distance between each degranulation event and the stimulatory surface (N ≥ 20 for each cell type). (D) RMA-s target cells were loaded with increasing concentrations of OVA and mixed with OT1 CTLs in the presence or absence of 30 μM nocodazole. Degranulation was measured by surface exposure of Lamp1. (E) OT1 CTLs expressing RFP and pHluorin-Lamp1 were imaged by confocal microscopy on glass surfaces coated with H2-K^b^-OVA and ICAM1 in the presence or absence of 30 μM nocodazole. The distance between each degranulation event and the stimulatory surface was measured and graphed (N ≥ 14 for each cell type). (F) RMA-s target cells were loaded with increasing concentrations of OVA and mixed with unpulsed RMA-s cells (bystanders) and OT1 CTLs at a 1:1:1 ratio either in the presence or absence of 30 μM nocodazole. Specific lysis of both targets and bystanders was assessed after 4h. In C-F, error bars denote SEM. P values (*, **, ***, and ns indicate P < 0.05, P < 0.01, P < 0.001, and not significant, respectively) were calculated by two-tailed Student’s T-test in all panels. All data are representative of at least two independent experiments.

IS formation is typically associated with the polarization of intracellular granules toward the interface. To quantify this process, we analyzed confocal images of CTLs on both stimulatory (H2-K^b^-OVA and ICAM1) and nonstimulatory (ICAM1 alone) surfaces. TCR engagement by H2-K^b^-OVA induced a marked shift of lytic granules toward the interface, which we detected by quantifying granzyme B (GzmB) staining intensity as a function of distance from the glass (Fig. S5A). Importantly, DKO CTLs polarized their granule pool in a similar manner, despite lacking centrioles. We also observed strong granule accumulation at DKO synapses by TEM (Fig. S5B). Hence, centrosomal integrity is dispensable for both granule polarization and synaptic secretion.

Both PCM proteins and the Golgi apparatus polarized to the IS in DKO CTLs (Fig. 1C, S1E), and it was therefore possible that these entities were providing enough microtubule organization to enable granule polarization and directional secretion in the absence of centrioles. To assess the importance of this residual organization, we treated wild type CTLs with nocodazole, a small molecule that induces microtubule depolymerization. Nocodazole abrogated the accumulation of lytic granules at the IS (Fig. S5C), implying an important role for microtubules in delivering granules to the cell surface for secretion. Consistent with this interpretation, overall levels of TCR-induced degranulation were markedly reduced in the presence of nocodazole (Fig. 4D). However, the directionality of granule release, as revealed by pHluorin-Lamp1 imaging, was unaffected. Indeed, although fewer degranulation events were observed in nocodazole-treated CTLs, those that did appear were polarized toward the stimulatory surface to the same extent as in untreated controls (Fig. 4E). We also assessed the effect of nocodazole on cytotoxic specificity by performing bystander killing assays.

Nocodazole treatment sharply reduced the lysis of OVA-loaded targets, consistent with its capacity to inhibit TCR-induced degranulation (Fig. 4F). Bystander killing was unaffected, however, remaining at the same low levels seen in control samples. We conclude that the microtubule cytoskeleton, while it does facilitate cytotoxic secretion, does not, in and of itself, control where lytic granule fusion occurs at the cell surface.

### Centrosome disruption impairs lytic granule formation

Because DKO CTLs did not exhibit a defect in directional secretion, we investigated alternative mechanisms that could explain their reduced killing potential. Cytotoxic capacity depends on the amount of perforin and granzyme available for use. Accordingly, we examined the expression of perforin and GzmB and found that DKO-CTLs contained markedly reduced levels of both proteins relative to wild type controls (Fig. 5A-B). This phenotype, which was evident by flow cytometry (Fig. 5A), immunoblot (Fig. 5B), and fluorescence imaging (Fig. S5A), manifested in DKO T cells as they differentiated into CTLs and lost their centrioles. At day 5 (three days after Cre transduction), DKO cells expressed normal amounts of GzmB, but by days 7 and 8 (five and six days after transduction, respectively) GzmB levels were approximately 2-fold lower than in wild type cells (Fig. 5A). *Sas4*^+/−^*Trp53*^−/−^ CTLs contained normal levels of GzmB (Fig. S2C), indicating that the DKO phenotype resulted from SAS4, and not P53, deficiency. DKO CTLs also released less GzmB after TCR stimulation (Fig. 5C), indicating that their diminished cytotoxic stores adversely affected their ability to mount secretory responses. DKO CTLs contained equivalent amounts of *Prf1* and *Gzmb* mRNA as controls (Fig. 5D), implying that their reduced expression of perforin and GzmB protein resulted from a post-transcriptional defect. We initially considered the possibility that, because of centrosomal dysfunction, DKO CTLs might spuriously release lytic granule contents in a TCR independent manner. If this were the case, one would expect to observe higher levels of cytotoxic factors in the culture medium. In fact, we observed more soluble GzmB in wild type CTL cultures (Fig. 5E), arguing against a model of inappropriate granule release.

**Figure 5.**
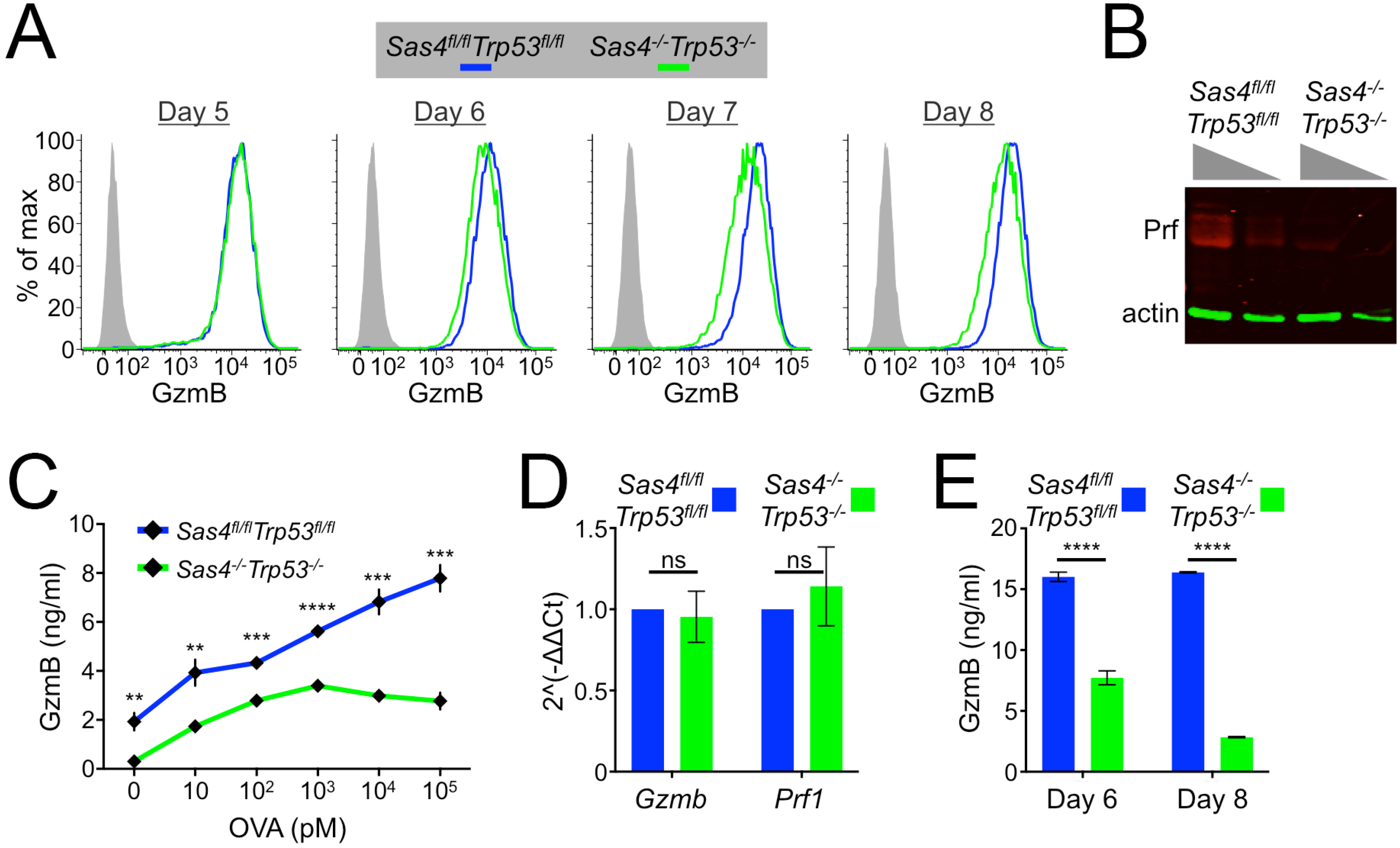
*Sas4*^*−/−*^*Trp53*^*−/−*^ CTLs carry and secrete less cytotoxic cargo. (A) *Sas4*^*fl/fl*^*Trp53*^*fl/fl*^ naïve OT1 T cells were stimulated with OVA-loaded splenocytes and then transduced with Cre expressing or control retrovirus after 48 hours. Subsequently, GzmB expression levels were assessed by flow cytometry in the differentiating CTLs at the indicated time points, which denote the number of days after initial antigen stimulation. (B) Perforin expression in day 8 *Sas4*^*fl/fl*^*Trp53*^*fl/fl*^ or *Sas4*^*−/−*^*Trp53*^*−/−*^ OT1 CTLs was evaluated by immunoblot, using actin as a loading control. Two dilutions of each sample were loaded, as indicated by the gray triangles. (C) RMA-s target cells were loaded with increasing concentrations of OVA and mixed with *Sas4*^*fl/fl*^*Trp53*^*fl/fl*^ or *Sas4^−/−^ Trp53^−/−^* OT1 CTLs. GzmB levels in the medium after 6 h were assessed by ELISA. (D) Quantitative RT-PCR analysis of *Gzmb* and *Prf1* mRNA expression in *Sas4*^*fl/fl*^*Trp53*^*fl/fl*^ or *Sas4*^*−/−*^*Trp53*^*−/−*^ OT1 CTLs. (E) GzmB levels in the culture medium of day 6 and day 8 *Sas4*^*fl/fl*^*Trp53*^*fl/fl*^ and *Sas4*^*−/−*^*Trp53*^*−/−*^ OT1 CTLs was quantified by ELISA. In C-E, error bars denote SEM. P values (**, ***, ****, and ns indicate P < 0.05, P < 0.001, P < 0.0001, and not significant, respectively) were calculated by two-tailed Student’s T-test in all panels. All data are representative of at least two independent experiments.

Reduced perforin and granzyme stores could also be caused by a defect in the maturation of lytic granules. To investigate this possibility, we stained wild type and DKO CTLs with lysotracker, a dye that labels lysosomal compartments. DKO CTLs exhibited substantially higher levels of lysotracker fluorescence as early as three days after Cre transduction (Fig. 6A), suggesting that centriole loss actually increases total lysosomal volume. We also assessed lysosome function by incubating CTLs with DQ-BSA, a fluorogenic protease substrate that is endocytosed readily by cells but only fluoresces upon degradation in lysosomes. DQ-BSA conversion was marginally slower in DKO CTLs than in wild type controls (Fig. S6A), a modest delay that was not caused by differences in endocytosis, as both populations robustly internalized TMR-BSA, a constitutively fluorescent substrate (Fig. S6A). Lysosomes also serve as a platform for signaling by the nutrient sensing mTORC1 complex. To assess mTORC1 function, we transferred nutrient-starved CTLs into complete medium and measured the phosphorylation of S6 kinase and 4EBP1, two canonical mTORC1 substrates. DKO CTLs exhibited markedly reduced mTORC1 activation (Fig. 6B), consistent with a defect in lysosomal identity or function. Collectively, these results indicate that centriole deletion alters both the volume and the activity of CTL lysosomes.

**Figure 6.**
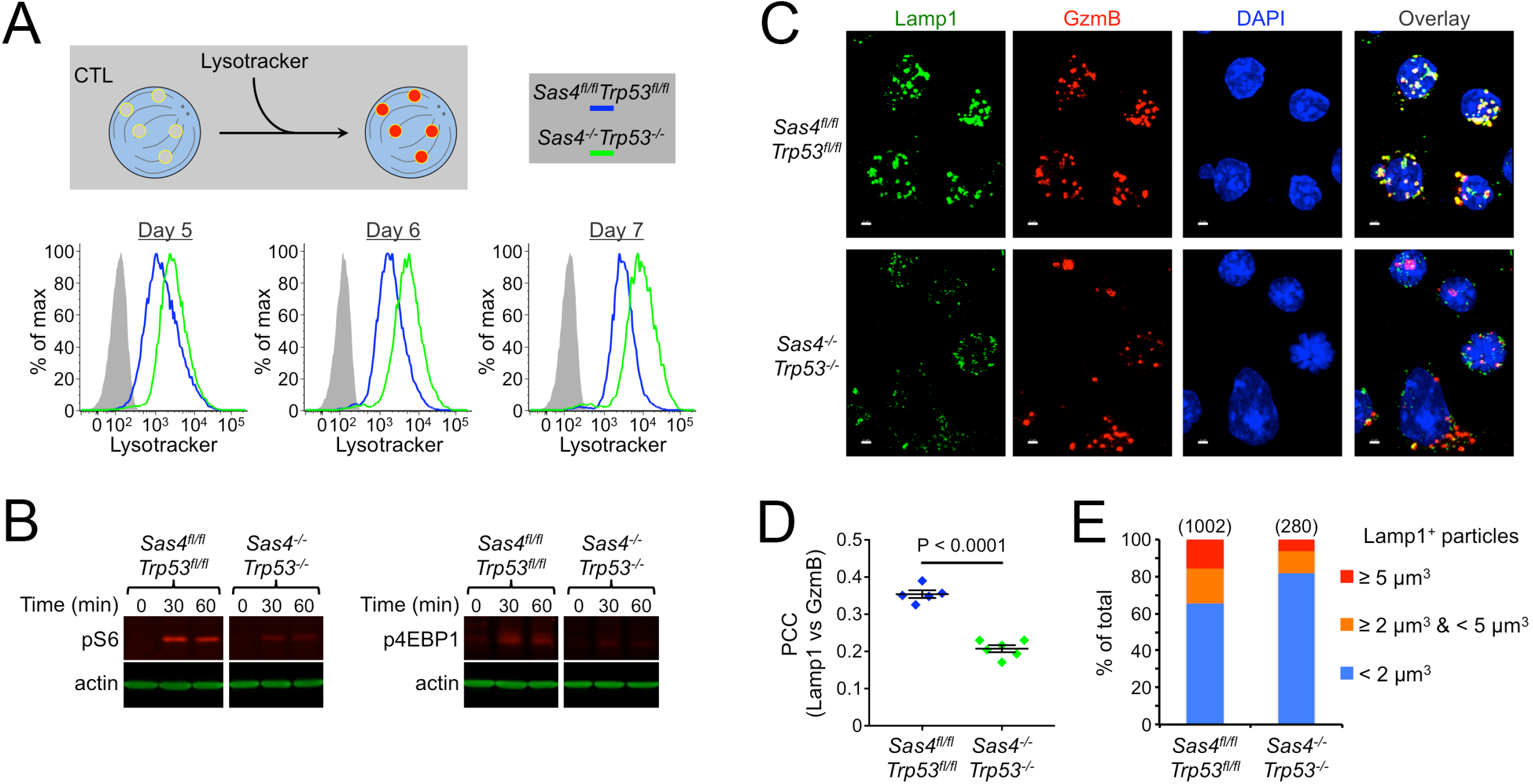
Centriole loss disrupts lytic granule biogenesis. (A) *Sas4*^*fl/fl*^*Trp53*^*fl/fl*^ naïve OT1 T cells were stimulated with OVA-loaded splenocytes and then transduced with Cre expressing or control retrovirus after 48 hours. Subsequently, lysosomal volume was assessed by lysotracker staining in the developing CTLs at the indicated time points, which denote the number of days after initial antigen stimulation. (B) Serum and nutrient starved *Sas4*^*fl/fl*^*Trp53*^*fl/fl*^ and *Sas4*^*−/−*^*Trp53*^*−/−*^ OT1 CTLs were incubated in complete medium for the indicated times, after which pS6 kinase and p4EBP1 were assessed by immunoblot, using actin as a loading control. (C) Representative confocal images of *Sas4*^*fl/fl*^*Trp53*^*fl/fl*^ and *Sas4*^*−/−*^*Trp53*^*−/−*^ OT1 CTLs stained with antibodies against Lamp1 and GzmB proteins. Nuclear DAPI staining is shown in blue. Scale bars = 2 μm. (D) PCC between Lamp1 and GzmB fluorescence was determined for each image of *Sas4*^*fl/fl*^*Trp53*^*fl/fl*^ and *Sas4*^*−/−*^*Trp53*^*−/−*^ OT1 CTLs (N ≥ 5). Each image contained 10-20 CTLs. P value was calculated by two-tailed Student’s T-test. (E) Graph showing the distribution of Lamp1^+^ particle size (see Methods) in *Sas4*^*fl/fl*^*Trp53*^*fl/fl*^ and *Sas4*^*−/−*^*Trp53*^*−/−*^ OT1 CTLs. The number of particles analyzed is indicated in parentheses above each bar. All data are representative of at least two independent experiments.

To further characterize this phenotype, we imaged CTLs stained for GzmB and the lysosomal protein Lamp1. Both molecules are established markers for lytic granules, and in wild type CTLs they exhibited strong colocalization, as expected (Fig. 6C). By contrast, many DKO CTLs contained cytoplasmic GzmB puncta with little to no associated Lamp1 staining (Fig. 6C). This colocalization defect was confirmed by Pearson’s Correlation Coefficient (PCC) analysis, which revealed significantly reduced overlap between Lamp1 and GzmB (Fig. 6D). DKO CTLs also contained a smaller fraction of large Lamp1 puncta, consistent with a defect in the accumulation of Lamp1 in lytic granules (Fig. 6E). Centriole loss did not alter the colocalization of Lamp1 with EEA1 and Rab7, which are markers for early and late endosomes, respectively (Fig. S6B-C), and it did not affect the size distribution of EEA1^+^ and Rab7^+^ compartments (Fig. S6B, S6D). These results implied that the granule organization phenotype seen in DKO CTLs did not result from the dysregulation of endosomal identity, but rather from a defect in the maturation of lysosomes into lytic granules. We conclude that centrioles are required for effective granule biogenesis.

### The centrosome regulates synaptic F-actin architecture and mechanics

The close spatiotemporal coupling between microtubule and F-actin dynamics at the IS raised the possibility centrosome disruption might alter F-actin remodeling. To investigate this hypothesis, we imaged OT1 CTLs by total internal reflection fluorescence microscopy on supported lipid bilayers containing H2-K^b^-OVA and ICAM1. Wild type CTLs form radially symmetric synapses with bilayers of this sort that are delimited by a peripheral, F-actin rich lamellipodium (Fig. 7A-B)(Le Floc’h et al., 2013; Sims et al., 2007). LFA1 accumulates at the inner aspect of this F-actin ring, forming a stereotyped “bulls-eye” structure. DKO CTLs exhibited a thicker, more irregular band of peripheral F-actin, which was accompanied by constriction of the LFA1 ring (Fig. 7B). To quantify this phenotype, we calculated the F-actin “clearance ratio”, a parameter that compares the fluorescence intensity at the edges of the IS with that at the center. A clearance ratio less than one is indicative of an annular fluorescence pattern, whereas a clearance ratio of one denotes a uniform distribution. This analysis confirmed that DKO CTLs were significantly impaired in their ability to form synaptic F-actin rings (Fig. 7C). We also carried out live imaging experiments using DKO or control CTLs that expressed the F-actin probe Lifeact-GFP. DKO CTLs often formed synapses that lacked an obvious F-actin cleared region at the center, and when these cleared regions did appear, they were more transient than in control CTLs (Fig. 7D). Collectively, these data indicated that centriole loss impairs proper F-actin remodeling at the IS.

**Figure 7.**
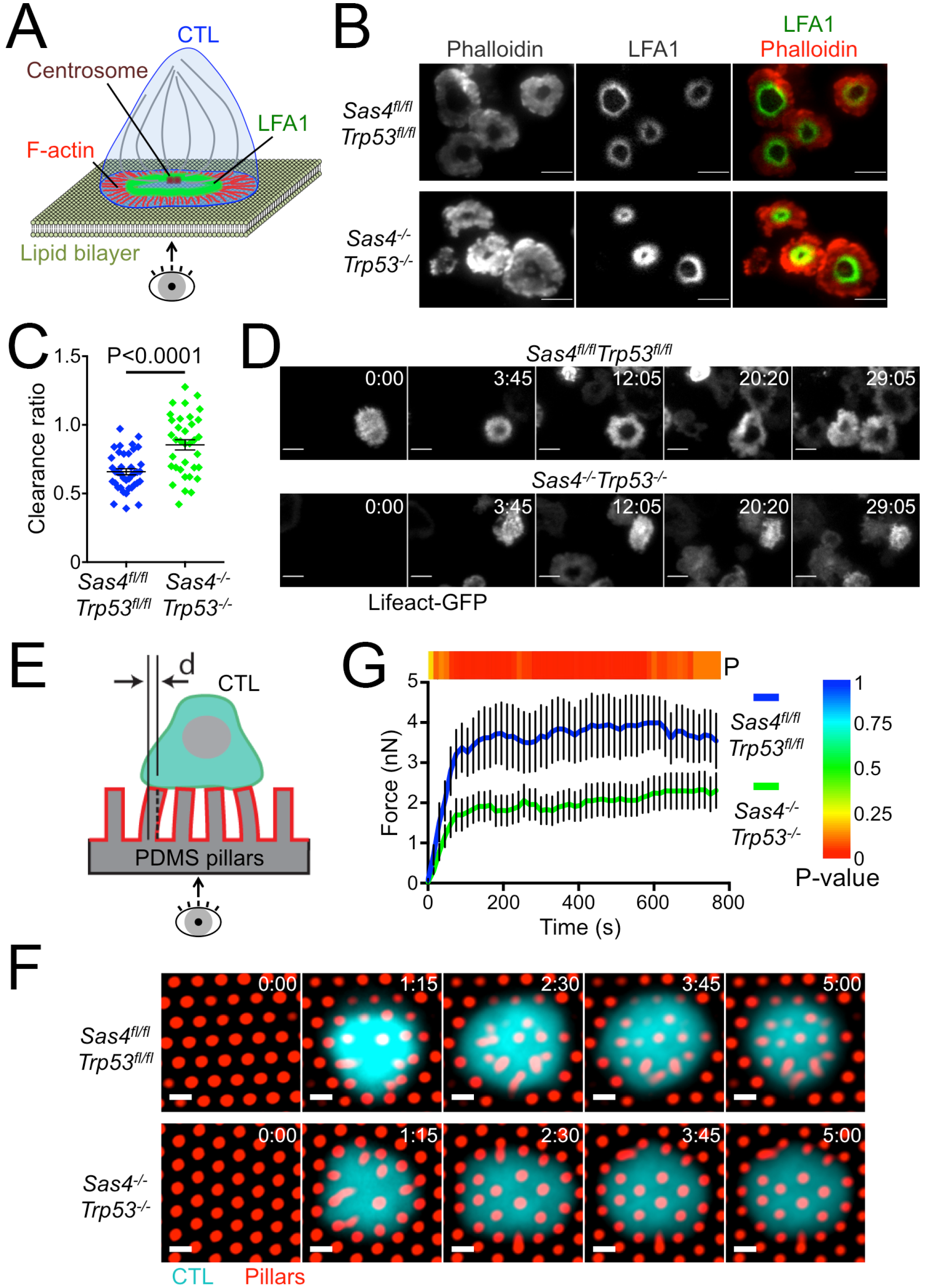
Centriole loss alters synaptic F-actin configuration and mechanical force exertion. (A) Schematic diagram of IS imaging on supported lipid bilayers. (B) *Sas4*^*fl/fl*^*Trp53*^*fl/fl*^ and *Sas4*^*−/−*^*Trp53*^*−/−*^ OT1 CTLs were added to supported lipid bilayers containing H2-K^b^-OVA and ICAM1, fixed, and stained with phalloidin (to visualize F-actin) and antibodies against LFA1. Representative TIRF images are shown. Scale bars = 10 μm. (C) F-actin clearance ratio (see Methods) was quantified for each cell (N ≥ 35 for each cell type). (D) Time lapse montages of representative *Sas4*^*fl/fl*^*Trp53*^*fl/fl*^ and *Sas4^-^/-Trp53^−/−^* OT1 CTLs forming synapses on supported lipid bilayers containing H2-K^b^-OVA and ICAM1. F-actin was visualized by TIRF imaging of transduced Lifeact-GFP. Time in MM:SS is indicated in the upper right corner of each image. Scale bars = 10 μm. (E) Schematic diagram of the micropillar system to measure synaptic forces. (F) *Sas4*^*fl/fl*^*Trp53*^*fl/fl*^ and *Sas4*^*−/−*^*Trp53*^*−/−*^ OT1 CTLs were labeled with fluorescent anti-CD45 F_ab_ fragment and added to PDMS micropillars coated with H2-K^b^-OVA and ICAM1. Time lapse montages of representative cells are shown. Time in M:SS is indicated in the upper right corner of each image. Scale bars = 2 μm. (G) Total force exertion against pillar arrays graphed versus time (N ≥ 11 of each cell type). Color bar above the graph indicates the P-value for each time point. In C and G, error bars denote SEM. P values were calculated by two-tailed Student’s T-test. All data are representative of at least two independent experiments.

Synaptic F-actin potentiates cytotoxicity by imparting mechanical force against the target cell (Basu et al., 2016). Accordingly, we investigated whether the F-actin disruption caused by centriole loss might compromise force exertion at the IS. OT1 CTLs were imaged on arrays of flexible polydimethylsiloxane (PDMS) micropillars that had been coated with H2-K^b^-OVA and ICAM1. T cells form IS like contacts with these arrays, inducing pillar deflections that can be translated into force vectors based on the known dimensions and composition of the pillars (Fig. 7E) (Bashour et al., 2014). DKO CTLs exerted significantly less force than their wild type counterparts (Fig. 7F-G), implying that centrosomal integrity is required for F-actin dependent mechanical output at the IS. Taken together with the data in the preceding section, these results indicate that centriole loss undermines killing capacity via two mechanisms, by 1) impairing lytic granule biogenesis, and 2) perturbing synaptic force exertion.

## Discussion

It is widely thought that centrosome reorientation guides directional lytic granule release at the IS (Liu and Huse, 2015; Stinchcombe and Griffiths, 2007). This model, which grew out of imaging studies of cytotoxic lymphocytes in the act of killing, elegantly explains both the potency and the specificity of the response. Lytic granules traffic along microtubules to access different intracellular locations. Thus, the idea that they reach the synaptic membrane by moving toward the centrosome in a minus end directed manner is eminently logical. Applying perturbation approaches to test this model, however, has been challenging because the molecular pathways that control centrosome reorientation also influence other critical aspects of lymphocyte activation. Here, we employed the alternative strategy of disrupting the CTL centrosome via centriole deletion. Although this approach altered microtubule architecture, it had no measurable effect on the polarization of lytic granules, the directionality of cytolytic secretion, and the specificity of target cell killing. It did, however, impair lytic granule biogenesis and synaptic F-actin dynamics. These results reveal an unexpected role for the centriole in controlling cytotoxic capacity and emphasize the importance of perturbation approaches when investigating complex cell biological mechanisms.

Although centriole deletion did perturb the organization of γ-tubulin and pericentrin, these PCM markers remained clustered in most DKO CTLs. TEM analysis of acentriolar cells revealed microtubules emanating from objects resembling PCM or centriole fragments, implying that this residual material retained microtubule-nucleating capacity. Clusters of residual PCM, together with associated microtubules, have also been observed in acentriolar DT40 B cells (Sir et al., 2013), further supporting the interpretation that they can mediate some degree of microtubule organization. We also note that the Golgi apparatus, which is known to nucleate and organize microtubules on its own (Efimov et al., 2007; Miller et al., 2009), retained its focal structure in the absence of centrioles. Importantly, both the Golgi and the residual PCM polarized toward the IS, which is one of the defining properties of the T cell centrosome. Hence, it seems likely that one or both of these structures served as a rudimentary MTOC in DKO CTLs, providing sufficient organization to guide the synaptic accumulation of lytic granules. This architectural redundancy is remarkable, and it highlights the importance of the microtubule cytoskeleton for T cell polarity and function.

The persistence of a rudimentary MTOC potentially explains how CTLs lacking centrioles remain capable of specific killing and directional granule release. It does not explain, however, the surprising functionality of CTLs lacking microtubules altogether. Indeed, while microtubule disassembly with nocodazole reduced the magnitude of degranulation responses, it failed to alter both cytotoxic specificity and directional secretion. Taken together, our results suggest that the microtubule cytoskeleton, while important for facilitating the efficient delivery granules to the cell surface, does not control where on the cell surface the granules fuse. That decision appears to be made by mechanisms that operate at the synaptic membrane itself.

What might these mechanisms be? Prior studies have demonstrated that degranulation occurs in plasma membrane domains within the IS that have been cleared of F-actin and are therefore accessible to fusion competent vesicles (Brown et al., 2011; Rak et al., 2011). The fact that granules do not fuse at actin hypodense regions outside the IS, however, implies that other factors also influence the process. Signaling lipids such as phosphatidylinositols and diacylglycerol are intriguing candidates in this regard. They control membrane specification, regulated secretion, and polarity in diverse cell types (Balla, 2013; Shewan et al., 2011), and they play a central role in the patterning of architectural microdomains within the IS (Gawden-Bone et al., 2018; Le Floc'h et al., 2013; Quann et al., 2009). Local Ca^2+^ influx, which drives regulated fusion at the neuronal synapse (Sudhof, 2012), could also be involved. Finally, recent studies have suggested that granule docking and fusion occur close to areas of strong integrin engagement (Houmadi et al., 2018), implying that outside-in signal transduction or mechanical tension could define permissive locations for exocytosis.

Certain cytokines, such as IL2 and IFNγ, are also secreted into the IS (Huse et al., 2006). Microtubules are strictly required for the directionality of this process, in stark contrast to what we have observed for lytic granules. In T cells, cytokines are trafficked directly from the Golgi apparatus to the cell surface along one of multiple directionally distinct constitutive pathways (Huse et al., 2006; Huse et al., 2008). Perforin and granzyme, by comparison, are stored for extended periods in acidic granules that inhibit their activity, and they are only released upon target cell engagement (Stinchcombe and Griffiths, 2007). Our present results make clear that T cells control the directionality of granule fusion via a microtubule independent mechanism at the cell cortex. This additional layer of regulation likely reflects the importance of restricting the effects of toxic perforin and granzyme to the target cell alone.

Although centriole deletion failed to disrupt directional lytic granule secretion in our hands, it did induce a marked and unexpected defect in granule biogenesis. DKO CTLs had increased lysosomal volume, but paradoxically exhibited delayed lysosomal degradation, reduced mTORC1 signaling capacity, and lower steady state levels of perforin and GzmB. They also exhibited reduced colocalization between Lamp1 and GzmB, implying a failure to sort cytotoxic proteins into fusion competent granules. Consistent with this interpretation, we found that these cells secreted less GzmB in response to TCR stimulation, despite normal surface exposure of Lamp1. Hence, not only do DKO CTLs store less perforin and granzyme, the factors that they do retain are also functionally handicapped due to improper compartmentalization. The microtubule cytoskeleton plays a well-established role in the intracellular trafficking of vesicular organelles. In recent years, however, it has become increasingly clear that microtubules are also critical for the maturation and function of these compartments. This is particularly true for lysosomes, whose degradative and signaling capacity depends on their location within the microtubule network (Korolchuk et al., 2011; Pu et al., 2016). Lysosome positioning is known to regulate cell migration and antigen presentation in immune cells (Alloatti et al., 2015; Bretou et al., 2017; Yuseff et al., 2011), and it will be interesting to investigate the extent to which centriole loss affects these processes.

Centrosome dysfunction has been linked to a number of inherited disorders, including polycystic kidney disease and Bardet-Biedl syndrome (Reiter and Leroux, 2017). The pathogenesis of these diseases, which are collectively called ciliopathies, has generally been attributed to defective primary cilia formation and function. Recent studies, however, have raised the possibility that certain symptoms characteristic of ciliopathies might be caused by cilia independent mechanisms, such as disrupted cell polarity and misoriented mitosis (Vertii et al., 2015). It is tempting to speculate that the dysregulation of compartmental identity of the sort observed in this study might also contribute to disease pathogenesis. In that regard, it is interesting to note that IFT20, and intraflagellar transport protein that is required for primary cilia formation and has been linked to ciliopathies, was recently shown to be required for proper lysosome biogenesis (Finetti et al., 2019).

The defects in synaptic architecture and force exertion observed in DKO CTLs imply an important role for the centrosome in shaping F-actin dynamics at the IS. It is known that the centrosome itself can directly nucleate F-actin (Farina et al., 2016), which could in principle function to organize the lamellipodial and protrusive structures emerging from peripheral IS domains. Alternatively, microtubules emanating from the centrosome could modulate cortical F-actin via proteins that bridge both cytoskeletal networks. In that regard, it is interesting to note that factors operating at the interface between F-actin and microtubules, such as the diaphanous formins and the scaffolding molecule IQGAP, have been implicated in centrosome reorientation and IS assembly (Andres-Delgado et al., 2012; Gomez et al., 2007; Gorman et al., 2012; Stinchcombe et al., 2006). Close coordination between the centrosome and F-actin dynamics could facilitate the efficient organization of synaptic output in space and time. Recent studies suggest that degranulation occurs in synaptic domains that are enriched in LFA1 and that are actively transmitting force against the target cell (Houmadi et al., 2018; Tamzalit et al., 2019). It will be interesting to investigate the role of the centrosome in ensuring this spatial coupling. Although the centrosome appears to be dispensable for directional secretion, it may yet be critical for linking granule fusion sites to adhesive or mechanically active zones within the IS.

In summary, our results demonstrate that an intact centrosome is indeed required for cellular cytotoxicity, but not for the reasons we initially hypothesized. In immune cells, organelles like the centrosome operate in the context of dramatic cytoskeletal plasticity, which can illuminate but also obscure their function. A comprehensive understanding of how these organelles contribute to immune cell biology will require not only close analysis of live imaging data but also experimental approaches that selectively disrupt components of interest. We anticipate that the continued application of targeted architectural perturbations like SAS4 deficiency will unearth additional unanticipated links between immune cell structure and function.

## Materials and methods

### Constructs

pMSCV vectors containing a linked cassette comprising a puromycin resistance gene, an internal ribosome entry site, and either CFP or RFP were prepared by subcloning the CFP and RFP coding sequences into pMSCV PIG using the NcoI and SalI restriction sites. The coding sequence for Cre recombinase was inserted into the resulting vectors (pMSCV PIC and pMSCV PIR) and pMSCV PIG using the BglII and EcoRI restriction sites in a process that destroyed the BglII recognition sequence. The retroviral pHluorin-Lamp1 and Lifeact-GFP constructs have been described (Le Floc’h et al., 2013; Rak et al., 2011).

### Cell culture and retroviral transduction

The animal protocols used for this study were approved by the Institutional Animal Care and Use Committee of Memorial Sloan-Kettering Cancer Center. *Sas4*^*fl/fl*^*Trp53*^*fl/fl*^ mice on the FVB background were crossed with OT1 TCR transgenic mice on the C57BL/6 background to obtain OT1 *Sas4*^*fl/fl*^*Trp53*^*fl/fl*^ and OT1 *Sas4*^*+/fl*^*Trp53*^*fl/fl*^ animals. To prepare primary CTL blasts, naïve T cells from these mice were mixed with irradiated C57BL/6 splenocytes pulsed with 100 nM OVA and cultured in RPMI medium containing 10% (vol/vol) FCS. Cells were supplemented with 30 IU/ml IL2 after 24 h and were split as needed in RPMI medium containing 10% (vol/vol) FCS and IL2. RMA-s cells were maintained in RPMI containing 10% (vol/vol) FCS. Phoenix E cells were maintained in DMEM medium containing 10% (vol/vol) FCS. For CTL transduction, Phoenix E cells were transfected with MSCV-Cre or empty MSCV together with packaging plasmids using the calcium phosphate method. Ecotropic viral supernatants were collected after 48 h at 37 °C and added to 1.5 × 10^6^ OT1 blasts 48 h after primary peptide stimulation. Mixtures were centrifuged at 1400 × g in the presence of polybrene (4 μg/ml) at 35 °C and the T cells then split 1:3 in RPMI medium containing 10% (vol/vol) FCS, 30 IU/ml IL2. 10 μg/ml puromycin was added to the cultures the following day to select for transduced cells. On day 5, transduced CTLs were further enriched by FACS and then cultured at 1.0 × 10^6^ cells/ml in RPMI containing 30 IU/ml IL2. Cultures of CTLs expressing Cre retrovirus were supplemented with the PLK4 inhibitor centrinone B (20 μM, MedChem Express) at day 5 in order to suppress the growth of any residual SAS4^+^ cells (Wong et al., 2015).

### Cytotoxicity experiments

RMA-s target cells were labeled with carboxyfluorescein succinimidyl ester (CFSE) or the membrane dye PKH26, loaded with different concentrations of OVA and mixed in a 96-well V-bottomed plate with CellTrace Violet (CTV) stained OT-1 cells. To assess killing, cells were mixed at a 1:1 or 1:10 E:T ratio and incubated for 4-24 h at 37 °C. In certain experiments, CTLs were combined with both OVA-loaded targets and unloaded bystander RMA-s cells at a 1:1:1 ratio. At the assay endpoint, specific lysis of RMA-s cells was determined by flow cytometry as previously described (Purbhoo et al., 2004). For degranulation experiments, the E:T ratio was 1:1 and cells were incubated at 37 °C for 90 min in the presence of eFluor 660-labeled Lamp1 Monoclonal Antibody (clone eBio1D4B, eBioscience). Lamp1 staining was then assessed by flow cytometry. In certain experiments, microtubules were depolymerized by preincubation for 10 min with 30 μM nocodazole, which was then maintained for the duration of the experiment. To measure conjugate formation, CTLs and targets were mixed 1:3, lightly centrifuged (100 × g) to encourage cell contact, and incubated for 20 min at 37 °C. Then, cells were then resuspended in the presence of 2% PFA, washed in FACS buffer (phosphate buffered saline (PBS) + 4% FCS), and analyzed by flow cytometry. Conjugate formation was quantified as (CFSE+ CTV+)/(CTV+).

### Western blots

To assess TCR-induced signaling pathways, serum and IL2 starved OT1 CTLs were incubated with streptavidin polystyrene beads (Spherotech) coated with H2-K^b^-OVA and ICAM1 at a 1:1 ratio for various times at 37 °C. To evaluate mTORC1 signaling, CTLs were starved in PBS and then stimulated for various times with RPMI containing 10% (vol/vol) FCS and IL2. Stimulated cells were lysed in 2 × cold lysis buffer (50 mM TrisHCl, 0.15 M NaCl, 1 mM EDTA, 1% NP-40 and 0.25% sodium deoxycholate) containing phosphatase inhibitors (1 mM NaF and 0.1 mM Na_3_VO_4_) and protease inhibitors (cOmplete mini cocktail, EDTA-free, Roche). Immunoblots were carried out using the following antibodies: pAkt (Phospho-Akt (Ser473) Ab; Cell Signaling Technology), pErk1/2 (Phospho-Thr202/ Tyr204; clone D13.14.4E; Cell Signaling Technology), IκB (Cell Signaling Technology), pS6 Kinase (Phospho-Thr389; clone 108D2; Cell Signaling Technology), and p4E-BP1 (Phospho-Thr389; clone 236B4; Cell Signaling Technology). Actin served as a loading control (clone AC-15, Sigma). Perforin levels were assessed by immunoblot using a polyclonal antibody (Cell Signaling Technology, #3693)

### Functional assays

To assess IFNγ and GzmB secretion, OT1 CTLs were mixed 1:1 with RMA-s cells loaded with increasing concentrations of OVA and incubated for 4-6 hours at 37 °C. Secreted IFNγ and GzmB were detected by ELISA using an established anti-IFNγ antibody pair (clone AN-18 for capture, eBioscience, biotinylated clone XMG1.2 for detection, BD Biosciences) or a mouse GzmB ELISA kit (Invitrogen #88-8022). In certain experiments GolgiPlug (BD Biosciences, manufacturers recommended concentration) was included in the assay medium and IFNγ production quantified by intracellular staining with fluorescently labeled anti-IFNγ antibodies (clone XMG1.2, TONBO). To assess TCR-induced proliferation, OT1 CTLs were CTV-labeled and mixed with OVA-loaded, irradiated C57BL/6 splenocytes. CTV dilution was assessed on a daily basis by flow cytometry. Intracellular GzmB levels were measured by flow cytometry using Alexa 647 labeled anti-GzmB (clone GRB11, Biolegend). To assess lysosomal content, CTLs were incubated with 50 nM lysotracker red (Thermofisher) for 60 min at 37 °C and then analyzed by flow cytometry. To assess lysosomal degradation, CTLs were mixed with 10 μg/ml DQ-BSA and 10 μg/ml TMR-BSA (both from Thermofisher) for various times at 37 °C, after which their green and red fluorescence was determined by flow cytometry. To monitor TCR downregulation, OT1 CTLs were mixed 1:1 with H2-K^b^- OVA- and ICAM1-coated beads at 37 °C for various times, after which TCR surface expression was assessed flow cytometrically using labeled antibodies against CD3ε (clone 500A2, eBioscience). Surface LFA1 was quantified using labeled antibodies against CD18 (clone M18/2, eBioscience).

### Quantitative RT-PCR

Total RNA from either control or DKO CTLs was extracted using the RNeasy Mini Kit (Qiagen), and RT-PCR was performed using the SuperScript™ III First-Strand Synthesis System kit (Invitrogen). Quantitative RT-PCR was performed using the Fast SYBR™ Green Master Mix (Applied Biosystems). Samples were processed using the BioRad CFX96 Real-Time System, and melting curve analysis was performed using BioRad-CFX Manager software. Gene expression level was determined by normalizing transcripts levels of the gene of interest to the level of the GAPDH housekeeping gene. The fold gene expression of each transcript relative to the control was calculated using the cycle threshold (Ct) method.

### Fixed imaging

Stimulatory glass surfaces and supported lipid bilayers bearing either H2-K^b^-OVA and ICAM1 or ICAM1 alone were prepared as described previously (Le Floc’h et al., 2013). OT1 CTLs were incubated for 10 min at 37 °C on these surfaces and then fixed by adding 4% paraformaldehyde for 5 min followed by ice-cold methanol for 15 min. Samples were then blocked in PBS with 0.5% Triton X-100 solution supplemented with 2% goat serum for 1 h at room temperature and incubated overnight at 4 °C with mixtures of the following primary antibodies: anti-centrin (clone 20H5; Millipore), anti-pericentrin (ab4448; Abcam), anti-γTubulin (clone GTU-88; Sigma), anti-βTubulin (clone TUB 2.1; Sigma), anti GM130 (clone 35/GM130; BD Biosciences), anti-Lamp1 (clone 1D4B; eBioscience), anti-GzmB (Polyclonal; ThermoFisher Scientific), anti-Rab7 (clone D95F2; Cell Signaling), anti-EEA1 (clone F.43.1; ThermoFisher Scientific). For F-actin staining, the methanol fixation step was omitted and cells were stained using Alexa 594-labeled phalloidin (ThermoFisher) and anti-LFA1 (clone M14/7; eBioscience). After primary antibody staining and washing, samples were incubated with the appropriate secondary antibody for 2 h at room temperature, washed and then imaged. In certain experiments, DAPI or Hoechst was added just prior to imaging to visualize cell nuclei. Confocal microscopy was performed with a Leica SP8 laser scanning microscope fitted with a white light laser and a 40 × objective lens. TIRF imaging was carried out using an Olympus OMAC system (IX-81 stage) outfitted with 561 nm and 488 nm lasers and a 60 × objective lens.

### Live imaging

For Ca^2+^ imaging, OT1 CTLs were loaded with 5 μg/ml Fura2-AM (ThermoFisher Scientific), washed, and then imaged on stimulatory glass surfaces coated with H2-K^b^-OVA and ICAM1. Images were acquired using 340 nm and 380 nm excitation every 30 seconds for 30 min with a 20 × objective lens (Olympus). For live imaging of degranulation, OT1 CTLs expressing pHluorin-Lamp1 and a fluorescent cellular label (typically RFP) were imaged on stimulatory glass surfaces coated with H2-K^b^-OVA and ICAM1 using a Leica Sp5 laser scanning confocal microscope outfitted with 561 nm and 488 nm excitation lasers and a 40 × objective lens. For each imaging run, a 12 μm z-stack (0.5 μm intervals) was collected every 15 s. In certain experiments, microtubules were depolymerized by preincubation for 10 min with 30 μM nocodazole, which was then maintained in the imaging medium for the duration of the experiment. To visualize synaptic F-actin dynamics, OT1 CTLs expressing Lifeact-GFP were added to bilayers containing H2-K^b^-OVA and ICAM1 and imaged every 5 s by TIRF microscopy for 30 min using a 60 × objective lens. For microwell cytotoxicity experiments, PDMS grids containing 50×50×25 μm wells were submerged in imaging medium and seeded with CTV-labeled OT1 CTLs and CFSE-labeled RMA-s cells that had been pulsed with 1 μM OVA. In general, individual wells contained between 1-3 CTLs and 1-3 RMA-s cells. 1 μM PI (Life Technologies) was included in the medium to enable real time visualization of dying cells. CTV-labeled CTLs were added and the cells imaged using a 20× objective lens (Olympus) at 10 min intervals for 12 hr. Brightfield, CFSE, CTV, and PI images were collected at each time point. The number of target cell killing events (identified by PI influx) was scored for each well and sorted based on the initial number of CTLs and targets in the well. Wells with no IS formation or with initial IS formation after 10 hours were excluded from the analysis.

### Electron microscopy

OT1 CTL populations were pelleted and resuspended in fresh, growth medium at 1×10^6^ cells/ml. Each CTL population was mixed 1:1 with EL4 targets pre-incubated for 40-60min with 1 μM OVA, left at room temperature for 5 min and incubated for 30-35 min at 37 °C to form conjugates. Conjugated suspensions were fixed by adding an equal volume of 5% gluteraldehyde/4% paraformaldehyde in PBS to give a final concentration of 2.5% gluteraldehyde/ 2% paraformaldehyde and left for 10 min, then the fixed cells were pelleted and fresh 1 × fixative in 0.1M Sodium Cacodylate added over the pellets. Pellets were subsequently processed for osmium fixation, urynal acetate staining en bloc and embedding in EPON as previously described for cells adhered on dishes (Jenkins et al., 2014; Tsun et al., 2011). 70-80 nm serial sections of CTLs conjugated to targets were collected on film-coated slot grids, stained with lead citrate, and the area of interest in each cell (identified by looking for the Golgi complex) followed over a depth of ~1-2 μm using an FEI Tecnai G2 Spirit BioTWIN transmission EM (Eindhoven, Netherlands). Images were captured using a Gatan US1000 CCD camera and FEI TIA software.

### Image analysis

Imaging data were analyzed using SlideBook (3I), Imaris (Bitplane), Excel (Microsoft), Prism (GraphPad), and Matlab (MathWorks). Ca^2+^ responses were quantified by first normalizing the ratiometric Fura2 response of each individual cell to the last time point before the initial influx of Ca^2+^, and then by aligning and averaging all responses in the data set based on this initial time of influx. Centrin^+^ puncta (Fig. 1B, S1B) were quantified by first establishing a high intensity threshold for all images of DKO and wild type control CTLs taken on the same day and then counting the number of fluorescent objects in each cell above this threshold. Radial analysis of microtubule and Golgi staining (Fig. 1C, S1D-E) was performed using a custom Matlab script. First, 180 linescans were taken through each cell at 1° increments and then averaged and amplitude-normalized to yield an intensity profile for that cell. Then, multiple intensity profiles (normalized for cell size) were averaged to generate the graphs shown in in the figures. Distances between degranulation events and the stimulatory surface (Fig. 4C, 4E) were determined with the Imaris Measurement Toolkit using yz or xz projections of confocal stacks. Pearson’s Correlation Coefficients (Fig. 6D, S6C) were calculated in Imaris for each image (10-20 cells per image). To determine the size distribution of intracellular compartments (Fig. 6E, S6D), 3-dimensional surfaces encompassing the compartments were first established in Imaris by intensity thresholding. Then, the volumes of all particles were output as an Excel file. For each compartment in question, identical intensity thresholds were applied to DKO and wild type control CTLs taken on the same day.

## Supplemental Figure Legends

**Figure S1.**
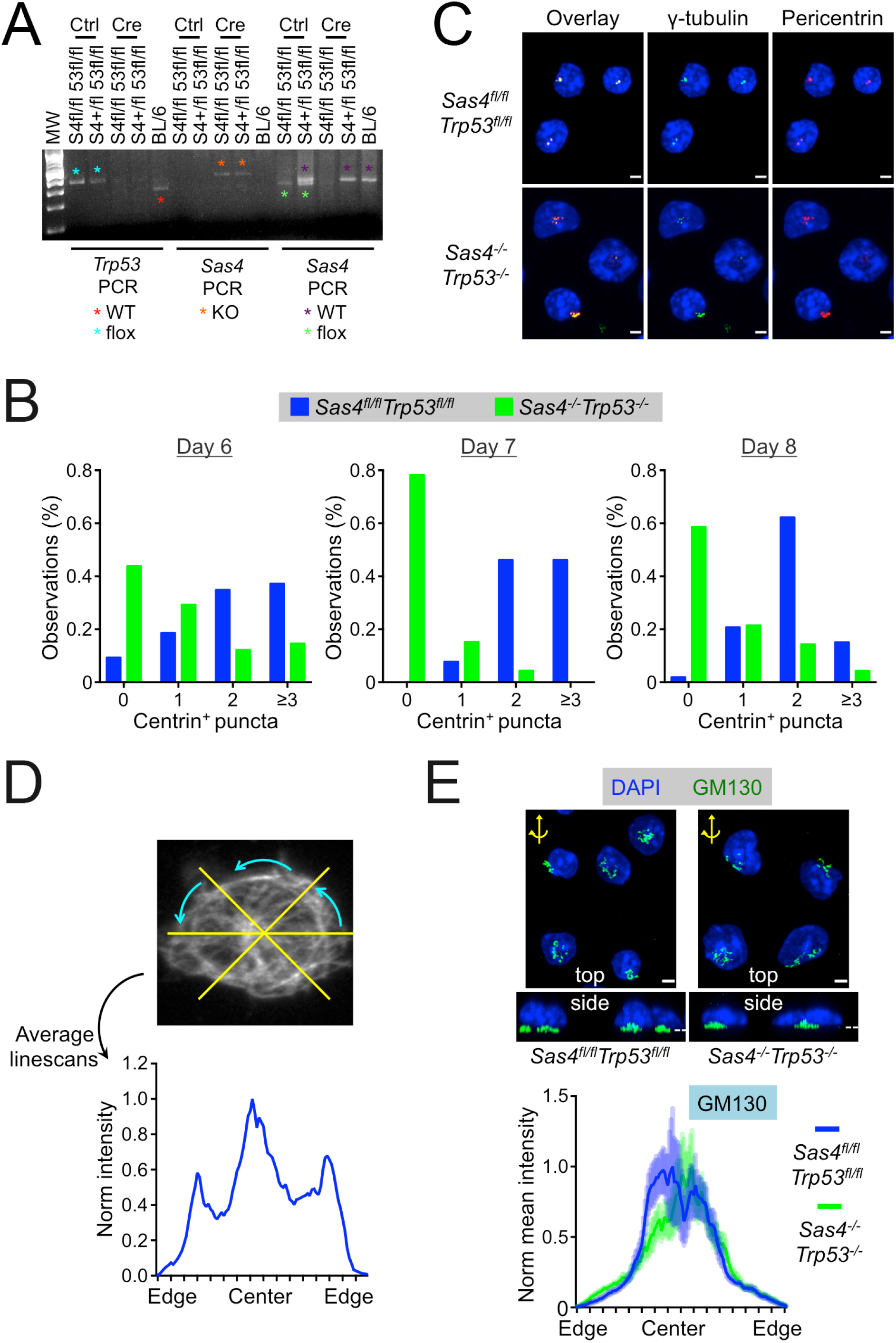
Architectural analysis of *Sas4*^*−/−*^*Trp53*^*−/−*^ CTLs. (A) PCR genotyping of the *Sas4 and Trp53* deletion. T cells from OT1 *Sas4*^*fl/fl*^*Trp53*^*fl/fl*^ (S3fl/fl 53fl/fl) and OT1 *Sas4^+/fl^Trp53^fl/fl^* (S4+/fl 53fl/fl) mice were stimulated with antigen and transduced with either control (Ctrl) or Cre-expressing retroviruses. After additional culturing and FACS purification, DNA extracts were PCR genotyped using primer mixtures for wild type and floxed *Trp53* (left), the *Sas4* knockout allele (middle), and wild type and floxed *Sas4*. Diagnostic bands are indicated with colored asterisks. Tail DNA from C57BL/6 mice (BL/6) was included as a wild type control. MW = molecular weight ladder. (B) *Sas4*^*fl/fl*^*Trp53*^*fl/fl*^ naïve OT1 T cells were stimulated with OVA-loaded splenocytes and then transduced with Cre expressing or control retrovirus after 48 hours. Subsequently, CTLs were stained with antibodies against centrin and the number of centrin^+^ puncta in each cell quantified at the indicated time points, which denote days after initial antigen stimulation. For each graph, N ≥ 39 *Sas4*^*fl/fl*^*Trp53*^*fl/fl*^ cells and N ≥ 41 *Sas4*^*−/−*^*Trp53*^*−/−*^ cells. (C) Representative confocal images of *Sas4*^*fl/fl*^*Trp53*^*fl/fl*^ and *Sas4*^*−/−*^*Trp53*^*−/−*^ OT1 CTLs stained with antibodies against the indicated centrosomal proteins. Scale bars = 3 μm. (D) Schematic diagram of the radial image analysis protocol. Multiple linescans (yellow lines on image to the left) were generated by rotating an initial horizontal line by 1° increments over 180°. These linescans were then averaged to yield an intensity profile for the entire cell (right), which was then averaged with other cell profiles to generate the curves shown in Figure 1C and Figure S1E. (E) Above, representative confocal images of *Sas4*^*fl/fl*^*Trp53*^*fl/fl*^ and *Sas4*^*−/−*^*Trp53*^*−/−*^ OT1 CTLs stained with antibodies against GM130. Top view (z-projection) images are shown above with corresponding side views (y-projections) below. Dotted white lines indicate the plane of the IS in the side views. The axis and rotation used to generate the side view is indicated in yellow in the top views. Scale bars = 2 μm. Below, normalized GM130 fluorescence intensity within radial domains between the center and the edge of the cell (N ≥ 12 for each cell type). Error bars denote standard error of the mean (SEM). All data are representative of at least two independent experiments.

**Figure S2.**
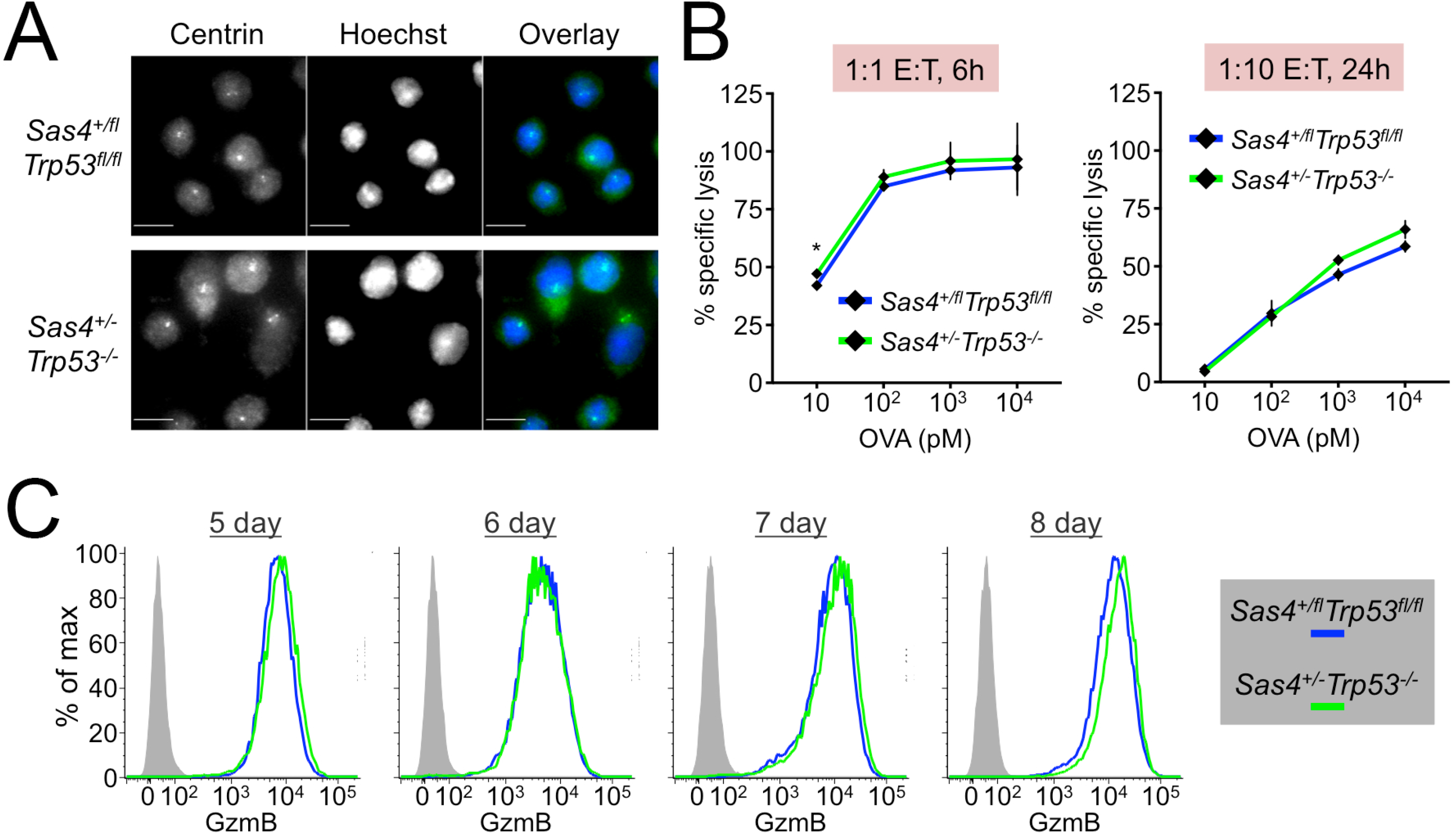
Architectural and functional analysis of *Sas4^+/−^Trp53^−/−^* CTLs. (A) Representative images of *Sas4^+/fl^Trp53^fl/fl^* and *Sas4^+/−^Trp53^−/−^* OT1 CTLs stained with antibodies against centrin. Scale bars = 10 μm. (B) RMA-s target cells were loaded with increasing concentrations of OVA and mixed with *Sas4^+/fl^Trp53^fl/fl^* or *Sas4^+/−^Trp53^−/−^* OT1 CTLs. Specific lysis of RMA-s cells was assessed at the indicated E:T ratios and times. (C) *Sas4^+/fl^Trp53^fl/fl^* naïve OT1 T cells were stimulated with OVA-loaded splenocytes and then transduced with Cre expressing or control retrovirus after 48 hours. Subsequently, GzmB expression levels were assessed by flow cytometry in the developing CTLs at the indicated time points, which denote the number of days after initial antigen stimulation. All data are representative of at least two independent experiments.

**Figure S3.**
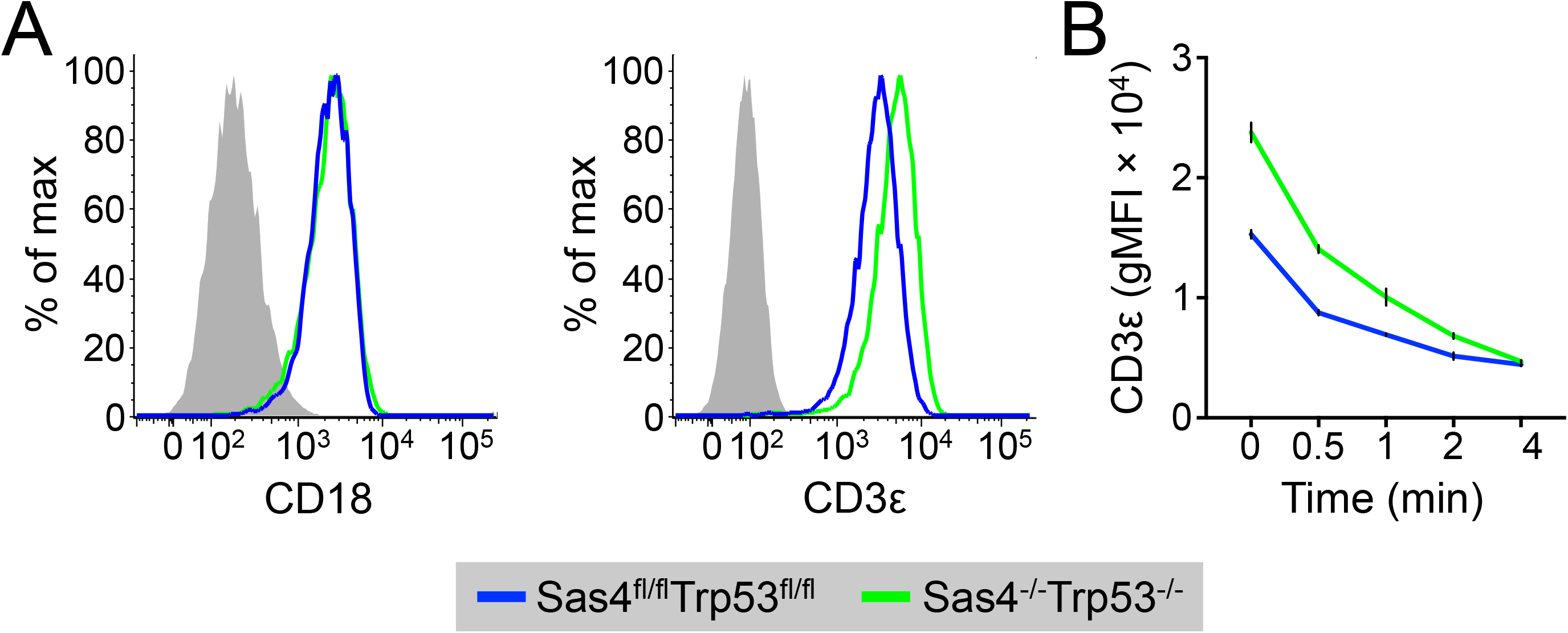
TCR and LFA1 expression of *Sas4*^*−/−*^*Trp53*^*−/−*^ CTLs. (A) Flow cytometric analysis of CD18 (LFA1 β chain) and CD3ε on *Sas4*^*fl/fl*^*Trp53*^*fl/fl*^ and *Sas4*^*−/−*^*Trp53*^*−/−*^ OT1 CTLs. (B) *Sas4*^*fl/fl*^*Trp53*^*fl/fl*^ and *Sas4*^*−/−*^*Trp53*^*−/−*^ OT1 CTLs were mixed with OVA-loaded RMA-s target cells and surface expression of CD3ε assessed by flow cytometry at the indicated times. Error bars denote SEM. Data are representative of at least two independent experiments.

**Figure S4.**
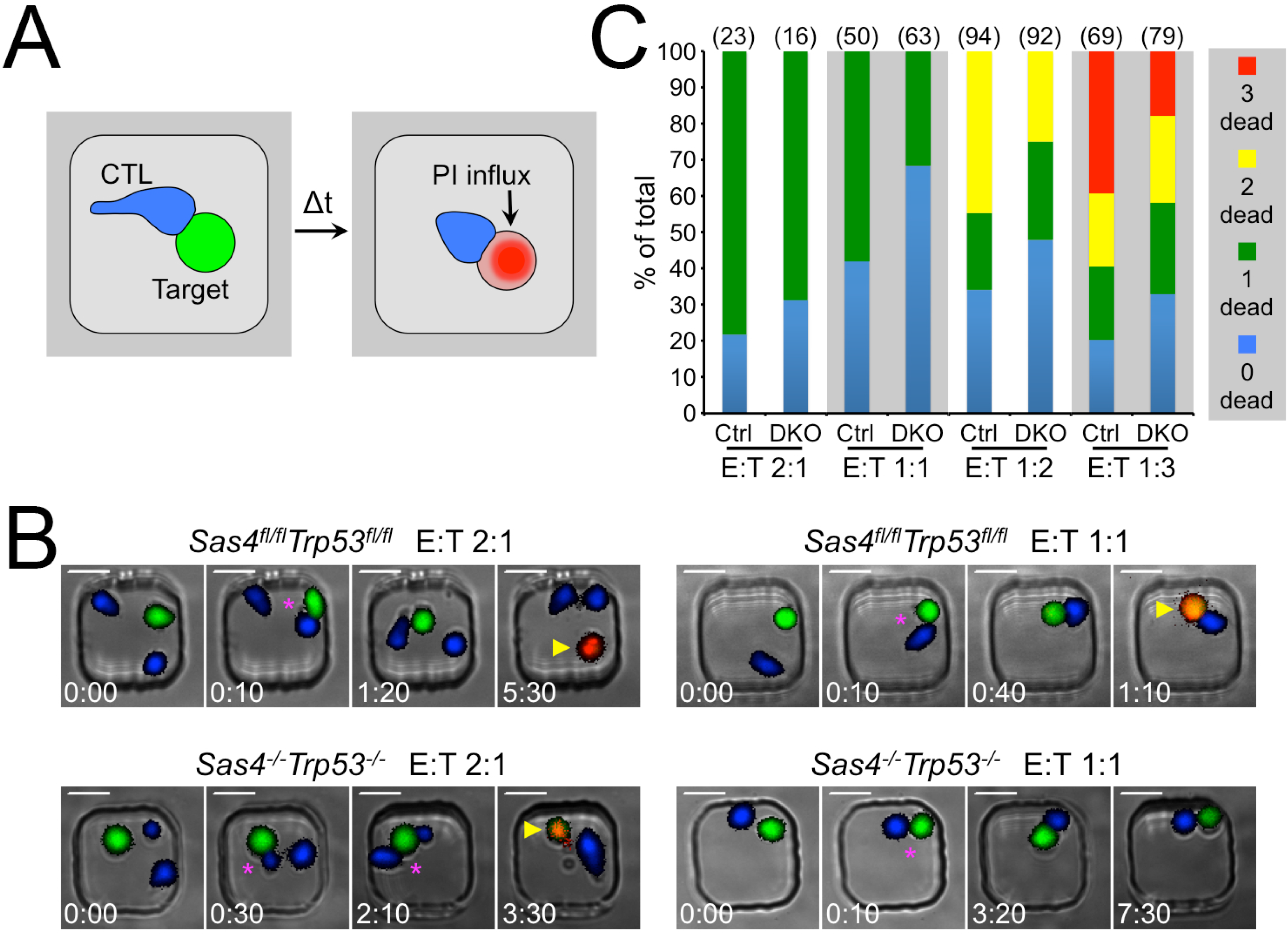
Imaging analysis of *Sas4*^*−/−*^*Trp53*^*−/−*^ CTL killing. *Sas4*^*fl/fl*^*Trp53*^*fl/fl*^ and *Sas4^−/−^ Trp53^−/−^* OT1 CTLs were mixed with OVA-loaded RMA-s target cells and loaded into 50 μm × 50 μm microwells to facilitate imaging of many CTL-target cell conjugates for 10-12 hours. (A) Schematic diagram showing detection of target cell death by PI influx. (B) Representative time-lapse montages of microwells containing one CFSE labeled target cell (green) and either one (right, E:T 1:1) or two (left, E:T 2:1) CTV-labeled CTLs (blue) of the indicated genotype. Magenta asterisks denote conjugate formation and yellow arrowheads the influx of PI. Time in H:MM is shown at the bottom left corner of each image. Scale bars = 10 μm. (C) Cumulative bar graph showing the distribution of outcomes in microwells containing the indicated E:T ratios. For each condition, the total number of wells analyzed is shown in parentheses above the bar. Ctrl = *Sas4*^*fl/fl*^*Trp53*^*fl/fl*^, DKO = *Sas4*^*−/−*^*Trp53*^*−/−*^. Data are representative of two independent experiments.

**Figure S5.**
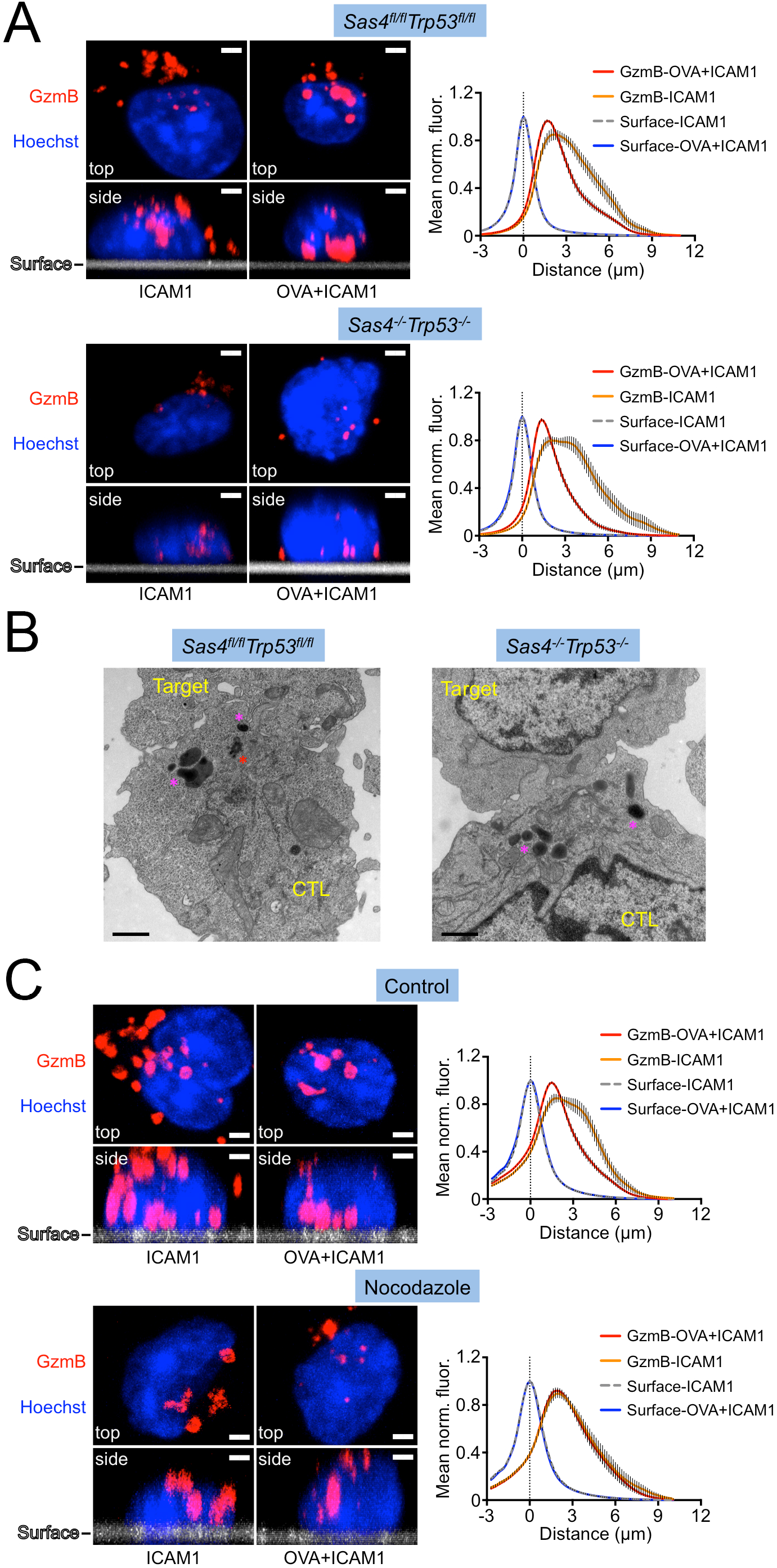
Centriole loss does not disrupt lytic granule polarization. (A) *Sas4*^*fl/fl*^*Trp53*^*fl/fl*^ and *Sas4*^*−/−*^*Trp53*^*−/−*^ OT1 CTLs were applied to glass surfaces coated with the indicated molecules (OVA = H2-K^b^-OVA) and then fixed and stained with antibodies against GzmB. Scale bars = 3 μm. Left, confocal images of representative CTLs shown from the top and from the side, with nuclear Hoechst staining is shown in blue. The stimulatory surface is indicated in side view images. Right, the average normalized fluorescence intensity distribution of the GzmB signal under each condition graphed as a function of distance from the stimulatory surface. The fluorescence distribution of the surface itself is also indicated. (B) TEM images of conjugates formed between *Sas4*^*fl/fl*^*Trp53*^*fl/fl*^ or *Sas4*^*−/−*^*Trp53*^*−/−*^ OT1 CTLs and OVA-loaded EL4 target cells. Magenta asterisks indicate lytic granules, and a red asterisk denotes the centriole in the *Sas4*^*fl/fl*^*Trp53*^*fl/fl*^ CTL. Scale bars = 1 μm. (C) OT1 CTLs were pretreated with 30 μM nocodazole or vehicle control (DMSO), applied to glass surfaces coated with the indicated molecules (OVA = H2-K^b^-OVA), and then fixed and stained with antibodies against GzmB. Left, confocal images of representative CTLs shown from the top and from the side, with nuclear Hoechst staining is shown in blue. The stimulatory surface is indicated in side view images. Scale bars = 2 μm. Right, the average normalized fluorescence intensity distribution of the GzmB signal under each condition graphed as a function of distance from the stimulatory surface. The fluorescence distribution of the surface itself is also indicated. All error bars denote SEM. Data are representative of two independent experiments.

**Figure S6.**
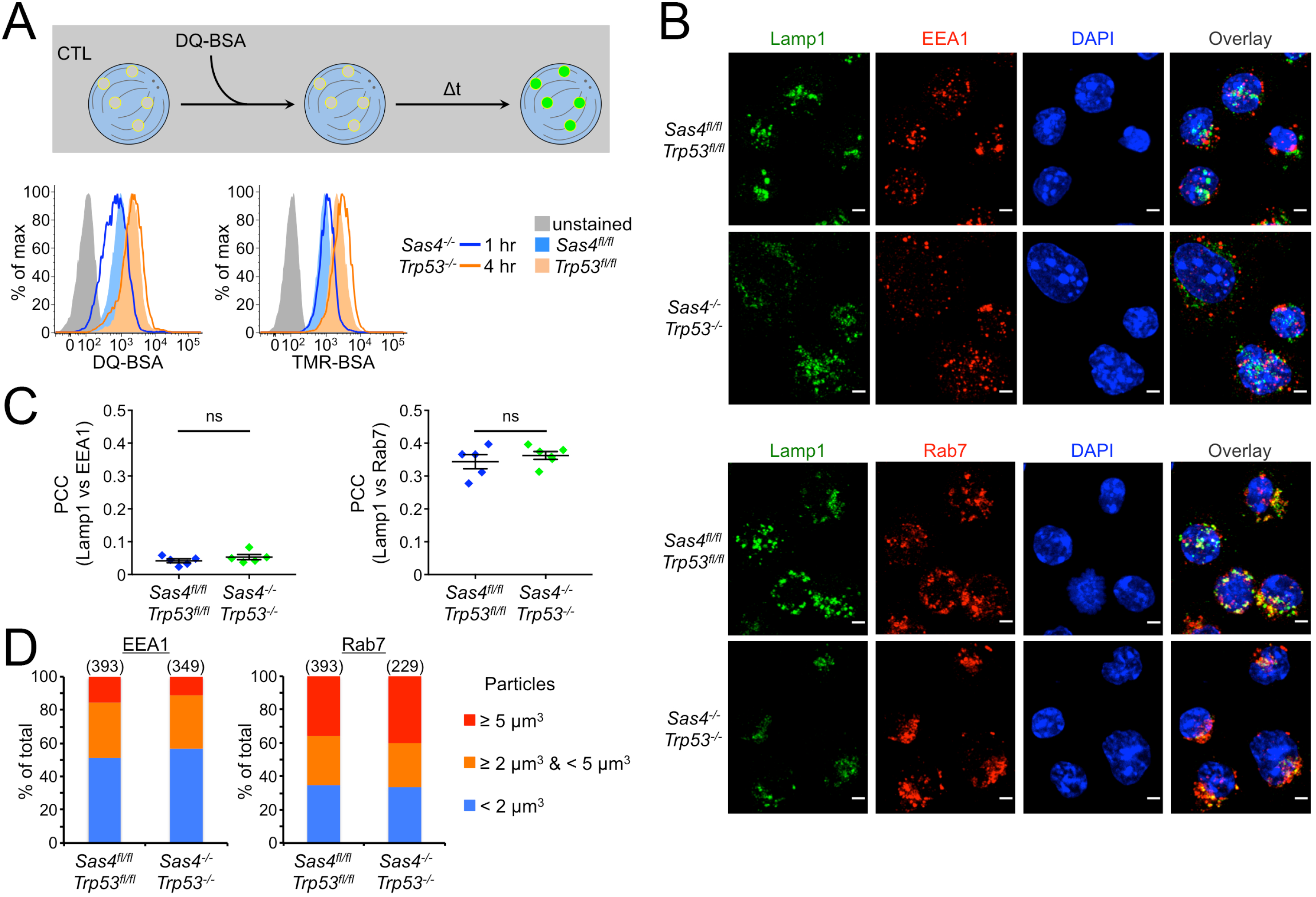
Endocytic and degradative capacity of *Sas4*^*−/−*^*Trp53*^*−/−*^ CTLs. (A) Above, schematic diagram of DQ-BSA uptake assay. Lysosomal degradation dequenches probe fluorescence. Below, flow cytometric analysis of DQ-BSA (left) and TMR-BSA (right) fluorescence at the indicated time points. (B) Representative confocal images of *Sas4*^*fl/fl*^*Trp53*^*fl/fl*^ and *Sas4*^*−/−*^*Trp53*^*−/−*^ OT1 CTLs stained with antibodies against the EEA1 and Lamp1 (above) and Rab7 and Lamp1 (below). Nuclear DAPI staining is shown in blue. Scale bars = 2 μm. (C) PCC between Lamp1 and EEA1 fluorescence (left) and between Lamp1 and Rab7 fluorescence (right) was determined for each image of *Sas4*^*fl/fl*^*Trp53*^*fl/fl*^ and *Sas4*^*−/−*^*Trp53*^*−/−*^ OT1 CTLs (N = 5). Each image contained 10-20 CTLs. P value was calculated by two-tailed Student’s T-test. (D) Graph showing the distribution of EEA1^+^ (left) and Rab7^+^ (right) particle size (see Methods) in *Sas4*^*fl/fl*^*Trp53*^*fl/fl*^ and *Sas4*^*−/−*^*Trp53*^*−/−*^ OT1 CTLs. The number of particles analyzed is indicated in parentheses above each bar. All data are representative of at least two independent experiments.

## Acknowledgments

We thank C. Firl for technical support; the MSKCC Molecular Cytology Core Facility for assistance with confocal imaging; the MSKCC Monoclonal Antibody Core Facility for fluorescently conjugated F_ab_; J. Stinchcombe and G. Griffiths (U. of Cambridge) for TEM analysis and critical reading of the manuscript; and J. Xavier (MSKCC), M. Overholtzer (MSKCC), Y. Lee (MSKCC), C. Lee (MSKCC), B. Tsou (MSKCC), M. Marks (U. of Pennsylvania), J. K. Burkhardt (U. of Pennsylvania), and members of the M. H. lab for advice. This work was supported in part by the NIH (R01-AI087644 to M. H., R01-AI110593 to L. C. K., P30-CA008748 to MSKCC) and the Cancer Research Institute (F. T.).

